# PEG-Free Tunable Poly(2-Oxazoline) Lipids Modulate LNP Biodistribution and Expression In Vivo after Intramuscular Administration

**DOI:** 10.1101/2025.06.05.657891

**Authors:** Colin Basham, Matt Haney, Yuling Zhao, Konstantin A. Lukyanov, Kyoungtea Kim, Ayse Baysal, Hallie Hutsell, Alexander V. Kabanov, Jacob D. Ramsey

## Abstract

Over the past decade, lipid nanoparticles (LNPs) have emerged as a transformative delivery platform, particularly in the field of mRNA vaccines, by enabling the stabilization and efficient intracellular delivery of nucleic acids. Importantly, the FDA approved LNPs used in the Pfizer-BioNTech and Moderna COVID-19 vaccines, rely on polyethylene glycol (PEG) to stabilize the nanoparticle *in vivo*. However, recent studies revealed that anti-PEG antibodies are ubiquitous in the population and are known to be a major cause of anaphylaxis and reduced therapeutic efficacy by accelerating blood clearance of PEGylated products. In this study, we report the development of novel poly(2-oxazoline) (POx) “stealth” lipids as PEG alternatives for LNP formulation. POx polymers are known to be poorly immunogenic and mitigate accelerated blood clearance. Upon varying polymer hydrophilicity and molecular weight, we screened transfection efficiency in multiple cell lines and evaluated top candidates *in vivo* by intramuscular administration. Our findings demonstrate that POx-lipids incorporating shorter hydrophilic methyl- and ethyl-oxazoline POx chains enhance transfection both *in vitro* and *in vivo*, while concurrently reducing liver accumulation and improving dendritic cell uptake - key features for effective vaccine delivery. These POx-lipids boost expression in muscle tissue by up to threefold, and greatly increase accumulation in extrahepatic tissues, including lymph nodes and spleen - critical sites for immune priming in the in the context of vaccination. Altogether, this work marks a notable advancement toward replacing PEG in LNP-based therapies and provides a clear path to optimizing POx-lipids for targeted vaccine and therapeutic applications.

## Introduction

Over the last decade, there has been a significant rise in nucleic acid therapies both under pre-clinical investigation and obtaining clinical approval^1,2^. The primary delivery vehicle for these therapies is the Lipid Nanoparticle (LNP), which encapsulates RNA in a lipid core, protecting it from degradation after administration and enabling intracellular uptake^3^. Prior nucleic acid delivery technologies suffered from high cost and poor *in vivo* transfection efficiency, precluding their translation to the clinic^4^. LNPs have enabled the intracellular delivery of many nucleic acid cargoes, with the most prominent example being the mRNA encoding for the viral spike protein during the COVID-19 pandemic. Despite their success, current LNPs have notable drawbacks including reactogenicity, inflammatory response, and immunogenicity, revealing new challenges in medicine^5,6^.

LNPs are comprised of four primary lipid components being the ionizable lipid, “helper lipid”, cholesterol, and pegylated (PEG, polyethylene glycol) lipid. The bulk of the formulation is typically the ionizable lipid and cholesterol, with approximately 2 mol % of the lipid being the pegylated lipid. “Pegylation” has long been used to stabilize nanoparticles *in vivo*, but recent studies have noted an increasing prevalence of anti-PEG antibodies in the general population since the COVID-19 pandemic^7,8^. While we have known that anti-PEG antibodies were a problem long before COVID-19, the pandemic and mRNA LNP vaccines have exacerbated the issue. One of the side effects of the LNP administration was anaphylaxis associated with anti-PEG immunity^9,10^. These responses were credited to the ubiquitous presence of PEG in commercial products ranging from pharmaceuticals to cosmetic products. This highlights the need for novel LNP technologies capable of evading immune detection while still producing stable particles. In addition, pegylated nanoparticles can induce accelerated blood clearance upon multiple administrations^7^. As a patient develops anti-PEG antibodies, the pharmacokinetic profile can become unpredictable and decrease overall exposure by accelerating the nanoparticles clearance from the bloodstream, significantly impacting therapeutic efficacy^10,11^. This highlights the needs for novel LNP technologies capable of evading immune detection while still producing stable particles.

Commercial LNPs tend to accumulate in the liver due to their interaction with apolipoprotein E (ApoE)^12,13^, causing hepatic toxicity, which is a concern for many treatments-particularly those requiring repeat dosing^11^. Most current work in the LNP field has focused on novel ionizable and helper lipid chemistries enabling alternative organotropism as researchers seek to inhibit liver accumulation and target other organs of interest^3,14^. For example, it is known that the addition of cationic phospholipids leads to preferential accumulation in the lungs while anionic modification leads to spleen accumulation^14^. In this work, we seek to modify the polymer lipid to achieve differential biodistribution and reduced immunogenicity of LNPs in the setting of intramuscular injections, which are commonplace for the administration of vaccines.

Our lab has worked to develop the Poly(2-Oxazoline) (POx) over the last two decades in the settings of small molecule, protein, and nucleic acid delivery^15–17^. We have demonstrated the low immunogenicity of our triblock copolymer micelles for the delivery of small molecule chemotherapies^18^. These triblock copolymers almost exclusively utilize the hydrophilic Poly(2-Methyl-2-n-Oxazoline) (PMeOx) as the corona forming block. This work will investigate the stability and transfection efficiency of LNPs formulated with non-PEG, POx polymers including PMeOx and Poly(2-Ethyl-2-n-Oxazoline) (PEtOx). Other labs have explored POx-based lipids for LNPs, with one study indicating PEtOx lipids mitigated accelerated blood clearance and reduced liver accumulation^19,20^. Yet, there has been limited work published with a systematic exploration of how POx polymer structure influences biodistribution and transfection efficacy -a key gap in the literature we seek to address in this work.

## Experimental

### Materials

#### Polymer Synthesis and Characterization

Methyl triflate was purchased from TCI Chemicals. Anhydrous acetontitrile, 2-n-2-methyl oxazoline, 2-n-2-ethyl oxazoline, sodium azide, deuterated chloroform, and calcium hydride were purchased from Sigma Aldrich. HPLC grade methanol, technical grade ether, HPLC grade DMF, and Lithium Bromide (LiBr) were purchased from Fisher Scientific. House deionized DI water was used for dialysis with 1, 2, or 3.5 kDa RC Spectrapor dialysis membranes used dialysis (purchased from Fisher Scientific). 0.22 µm nylon syringe filters (13 mm) were purchased from Thermofisher. Poly Methyl Methacrylate (PMMA) Gel Permeation Chromatography (GPC) standards were purchased from Sigma Aldrich.

#### Lipid Synthesis

16:0 DBCO-PE (1,2-dipalmitoyl-sn-glycero-3-phosphoethanolamine-N-dibenzocyclooctyl) lipid was purchased from Avanti Research and 200-proof anhydrous ethanol (Supelco) was purchased from Sigma Aldrich.

#### LNP Formulation and Characterization

Cholesterol, DSPC (1,2-distearoyl-sn-glycero-3-phosphocholine), 16:0 PEG-2000 PE (1,2-dipalmitoyl-sn-glycero-3-phosphoethanolamine-N-[methoxy(polyethyleneglycol)-2000] (ammonium salt)) were purchased from Avanti Research, while ionizable lipid SM-102 was purchased from Broadpharm. Chloroform was purchased from Fisher Scientific and 200-proof anhydrous ethanol (Supelco) was purchased from Sigma Aldrich. Sodium Acetate and Glacial Acetic Acid were purchased from Sigma Aldrich. Firefly luciferase mRNA was purchased from TriLink Bio. UltraPure DNAase/RNAase free water (Invitrogen) was purchased from Fisher Scientific and 1X DPBS (Gibco) was purchased from UNC Tissue Culture Facility. Sterilized House DI water was utilized. Quant-it Ribogreen reagent was purchased from ThermoFisher Scientific. DPBS used for size, particle concentration, and zeta potential analysis was filtered with 0.2 µm nylon syringe filters (25 mm) from Fisher Scientific. Cuvettes for dynamic light scattering (DLS) (Product ZEN0040) were purchased from Malvern.

#### *In Vitro* Transfections

HEK293 cell line was purchased form ATCC, and DC2.4 cell line was donated in kind from the Vincent lab at UNC Chapel Hill (previously purchased from Sigma Alrich). 4T1 cells were previously given in-kind from the Perou Lab (UNC Chapel Hill). RPMI and DMEM media (Gibco) cell culture medium and Avantor Fetal Bovine Serum (FBS) were purchased from the UNC Tissue Culture Facility. Triton X-100 reagent was purchased form Fisher Scientific. Luciferase Assay Kit was purchased from Promega. BCA Protein Assay Kit (Pierce) was purchased from Fisher Scientific. Antibiotic/Antimycotic, L-glutamine, MEM Non-essential amino acids, and 2-mercaptoethanol were purchased from Gibco.

#### *In Vivo* Transfections

Mice (Balb/c) were purchased from Jackson Labs. D-Luciferin potassium salt for *in vivo* use from Biosynth. Triton X-100 reagent was purchased from Fisher Scientific. Luciferin Assay Kit was purchased from Promega for analysis of *ex vivo* organs.

#### Instrumentation

Mettler Toledo XS105 Dual Range Analytical Balance. Nicolet 380 (ThermoFisher Scientific, Waltham, MA) FTIR. Malvern OMNISEC GPC/SEC. Bruker AVANCE NEO 400 MHz Spectrometer ^1^HNMR. Malvern Zetasizer Nano ZS (Malvern Panalytical Ltd., UK). Zetaview QUATT Nanoparticle Tracking Analyzer PMX-420 (Particle Metrix, Germany). Nanodrop 2000C Spectrophotometer (ThermoFisher). Molecular Devices Spectramax M5 Plate Reader. PROMEGA 20/20 GLOMAX Luminometer. IVIS Spectrum (Revvity Inc.) *in vivo* Imaging System. Helix Bio Impinged Jet Mixer.

### Polymer Synthesis

POx polymers were synthesized by living cationic ring opening polymerization (LCROP). Briefly, all monomers were distilled on a schlenk line under reduced pressure/argon atmosphere to purify and remove water. Water content was confirmed below 200 ppm by Karl Fisher titration prior to use in synthesis. All glassware was dried at 120 °C overnight before use. Glassware was allowed to come to temperature under argon flow. Anhydrous acetonitrile was added to the dry schlenk flask followed by desired amount of initiator (methyl triflate). Monomer was then added to the reaction flask to achieve the desired degree of polymerization. The schlenk flask was sealed and heated at 100 °C overnight. Reaction progress was monitored by ^1^HNMR and reaction was allowed to proceed until ∼100% monomer conversion. After full conversion and desired degree of polymerization (DP) was reached, polymers were terminated with 10-fold molar excess sodium azide and allowed to react overnight at 100 °C. After termination, the reaction mixture was dissolved in small amount of methanol and centrifuged at 2,000 x G for 5 minutes to pellet any excess, undissolved sodium azide. Solvent was removed under reduced pressure and the polymer was dissolved again in minimal methanol. It was again centrifuged at 2,000 x G for 5 minutes to remove excess sodium azide. The polymer-methanol solution was then precipitated in ∼30-fold volumetric excess ice-cold diethyl ether under rapid stirring. Ether was decanted. Polymers with DP 10 were dissolved in minimal methanol and precipitated once more in ether. DP 10 polymers were then dissolved in DI water and lyophilized. Polymers greater than DP 10 were dissolved in DI water and dialyzed in 1 kDA dialysis membrane (DP=20), 2 kDa dialysis membrane (DP=40), or a 3.5 kDa membrane (DP=80). Polymers were dialyzed against 5 L of DI water for 3 days, switching out water daily prior to lyophilization.

### ^1^HNMR Characterization

^1^H NMR spectrum was acquired using a Bruker AVANCE NEO 400 MHz Spectrometer and analyzed using the MestReNova software. Spectra were calibrated using residual solvent signal. Number-average molecular masses (M_n_) were determined via end-group analysis using ratio of initiator methyl protons to backbone and side chain protons after purification.

### GPC Characterization

GPC was performed to determine polymer molecular weight and molecular weight dispersity with a Malvern OMNISEC GPC/SEC system using a DMF (N,N-Dimethylformamide, 0.1% LiBr) mobile phase and two PolarGel-L columns (PL1117-6830) in sequence. Samples were dissolved overnight in the mobile phase at 5 mg/mL, then passed through a 13 mm 0.22 μm nylon filter into autosampler vials. Samples were analyzed with a refractive index detector during a 30 minute method with a 0.8 mL/min flow rate, 100 μL injection volume, and column and detector temperature of 25°C. A 3^rd^ order polynomial fit was used to evaluate the data using conventional calibration with PMMA standards ranging from 2,000 to 20,000 g/mol.

### FTIR Characterization

The azide termination of POx polymers was verified using single bounce Attenuated Total Reflectance Fourier Transform Infrared spectroscopy (ATR-FTIR). Using a Nicolet 380 (ThermoFisher Scientific, Waltham, MA) FTIR solid polymers were pressed onto a diamond crystal surface and the attenuated total reflectance of an infrared laser interacting with the sample was recorded from 400 cm^-1^ to 4000 cm^-1^. Spectra were averaged across 2000 runs at 4 cm^-1^ resolution and adjusted using advanced ATR correction and baseline correction in the OMNIC software. The presence of a characteristic azide peak around 2100 cm^-1^ was confirmed after purification of each polymer.

### Partial Specific Volume Measurements

The partial specific volume of each polymer was determined using a 1 mL glass pycnometer. The pycnometer was first calibrated by filling with water to determine the exact volume. Each polymer studied in this work was then prepared as a 25 mg/mL solution in water. The glass pycnometer was filled with each solution and the weight measured. For each measurement, the vial was weighed six times to obtain an average weight. Care was taken to ensure the vessel was filled precisely to the calibration mark with no air bubbles or overfill. The polymer solution was weighed, and the amount of water in the solution vial was determined by subtracting the total weight of the solution from the known amount of polymer which was added to the pycnometer. Using the density of water (0.997 g/mL at 25 °C), the volume of polymer was obtained and the partial specific volume of said polymer calculated.

### Lipid Conjugation to Polymers

Lipids were conjugated to polymers using strain-promoted alkyne-azide cyclo-additon (SPAAC) click chemistry. 16:0 DBCO-PE phospholipids were dissolved in ethanol at 1 mg/mL, and azide-terminated POx polymers were dissolved in ethanol at 3X molar excess. UV absorbance at 310 nm was monitored on a Nanodrop 2000C Spectrophotometer (ThermoFisher). The sample was vortexed and stored overnight at 4 °C. Successful click conjugation was confirmed through the disappearance of the UV absorbance peak at 310 nm which characteristic of unreacted DBCO.

### LNP Preparation

Four-component LNPs were prepared containing SM-102 ionizable lipid, cholesterol, DSPC, and PEG- or POx-ylated lipid in a 50:38.5:10:1.5 mol% ratio. Briefly, lipids stored in chloroform were combined and placed in a vacuum chamber until the solvent was fully evaporated. Then, the dry lipid film was solvated with ethanol to the requisite volume for mixing. Firefly luciferase (fLuc) mRNA was thawed at 4 °C and dissolved in a 10 mM acetate buffer at pH 4.15.

Lipids in ethanol and mRNA in buffer were combined at an amine/phosphate (N:P) ratio of 5.2:1 using a Nova Benchtop impinged jet mixer (Helix Biotech), using a flow rate ratio of 3:1 and a total flow rate of 8 mL/min. Formulated LNPs were immediately placed in dialysis tubing (10 or 12 – 14 kDa MWCO) and dialyzed against 10 mM acetate buffer, pH 4.15 for 4 – 6 hours at 4 °C; then, dialysis buffer was replaced with 1X DPBS and LNPs were dialyzed overnight at 4 °C. LNPs were collected and concentrated using Amicon Ultra-4 centrifugal filter units (100 kDa MWCO).

### DLS Size and Polydispersity Analysis

Concentrated LNPs were diluted 100X into 1X DPBS. They were then diluted 5X into DPBS for size and polydispersity analysis with a Malvern Zetasizer Nano. Separate aliquots were diluted into 1X DPBS buffer, and particle size and polydispersity were characterized on a Malvern Zetasizer Nano ZS. Three separate measurements were made for each prepared batch and averaged together to obtain Intensity Size, Number size, Z-Average Size, and PDI.

### Zetaview Size, Concentration, and Zeta Potential Analysis

After an initial 100X dilution in DPBS, particles were further diluted 1,000X into 0.2 μm filtered DPBS (Size and Concentration analysis) or 0.2 μm filtered 0.1X DPBS (Zeta Potential Analysis to reduce solution conductivity). For size and concentration analysis, two measurements were obtained at each of 11 positions in the sample cell. For Zeta Potential, a measurement was performed at every position. We report mean, median, and mode size, particle concentrations, and zeta potential.

### Ribogreen RNA Encapsulation Assay

The encapsulation efficiency and mRNA concentration was measured using a modified version of the Quant-it Ribogreen Reagent protocol. After an initial 100X dilution in DPBS, LNPs were further diluted 20X into either 1X Triton to lyse or in 1X TE Buffer supplied with the Ribogreen kit. They were each allowed to sit for 1 hour at room temperature before assaying. A 2000X dilution of the Ribogreen reagent was prepared in TE Buffer (need 100 uL per sample well. All samples and standards were run in duplicate). The standard RNA was diluted to 2 ug/mL as described in the protocol, and we confirmed the concentration by measuring the absorbance at 260 nm via nanodrop. A standard curve ranging from 2-100 ng RNA/mL was prepared. After 1 hour, each sample was further diluted 5-fold into TE buffer. 100 uL of sample or standard was added in duplicate to a black 96-well plate followed by 100 uL of the 2000X diluted working reagent. A blank TE buffer control was also used as a sample for background subtraction. After 30 minutes, the fluorescence was read on a Molecular Devices SpectraMax M5 plate reader with 480 nm excitation and 520 nm emission. The encapsulation efficiency was calculated according to **Equation 1** below.

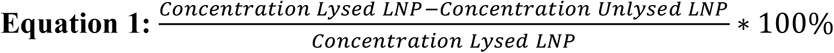

### *In Vitro* Transfection

HEK293, DC2.4, or 4T1 cells were seeded in appropriate medium at 30,000 cells/well in a 96-well plate with n=6 wells prepared for each treatment group including blank and mRNA alone controls. HEK293 was cultured in DMEM with 10% FBS and 1% antibiotic/antimycotic. DC2.4 were cultured in MEM media with 10% FBS, 1% antibiotic/antimycotic, 1X L-glutamine reagent, and 0.0054X 2-mercaptoethanol. 4T1 cells were cultured in RPMI media with 10% FBS and 1% antibiotic/antimycotic. Cells were allowed to adhere overnight. LNPs were diluted in appropriate cell culture media to 1 μg/mL total mRNA. For treatment, media was removed and 100 μL of LNPs were added to each well (100 ng mRNA/well). Cells were incubated for 24 hours. At 24 hours, media was removed and cells were lysed in 30 μL of 1X Triton and incubated with shaking at 4 °C for 1 hour. After one hour, 2 μL of each well was pipetted into a microcentrifuge tube (transparent). Sequentially and read one at a time, 50 μL of Luciferin assay reagent was added to each tube and luminescence read immediately on a ProMega GLOMAX 20/20 Luminometer. After luminescence readings, cell lysate was diluted 10X in 100-fold diluted Triton X-100. Cell lysates were then assayed according to standard BCA assay protocol for protein concentration while diluting standards in 1X Triton. Luminescence was normalized to the μg of protein in the lysate. Each individually prepared LNP batch was assayed *in vitro*.

### *In Vivo* Transfection

For *in vivo* studies, the two batches of each formulation were combined into a single aliquot for administering to mice with equal mRNA contributions coming from each one. 325 Balb/c mice (3 female, 3 male) were injected intramuscularly with 5 μg of fLuc mRNA in a 50 μL volume (DPBS diluent). Prior to injections, mice were shaved so that fur would not interfere with luminescence signals. After 4 hours, mice were injected intraperitoneally with 50 μL of 30 mg/mL D-Luciferin *in vivo* reagent. After 5 minutes, they were put under isoflurane anesthesia, and 5 minutes later luminescence was imaged on an IVIS Spectrum *in vivo* Imaging System. Images were acquired of side views and ventral views ensuring no saturated pixels were present in an image. Regions of interest (ROIs) were drawn around muscle tissue (side view) and livers (ventral view) for quantification of mRNA expression. Imaging was repeated at 24 hours post-injection.

### Ex Vivo Organ Luminescence Analysis

After the 24 hour imaging, mice were euthanized and muscle tissue (site of injection), liver, kidney, spleen, draining lymph nodes, lung, heart, and brain were collected and small chunks were weighed into previously weight microcentrifuge tubes so the weight of the tissue was known. 100 μL of 1X Triton was added to each tissue (except lymph nodes where 50 μL was added) and the tissues were frozen at −80 °C overnight. In the morning, tissues were homogenized with plastic pestle which fits into 1.5 mL microcentrifuge tubes. Pestle was rinsed with water and dried between tissue samples. After homogenization, the samples were left at 4 °C for two hours before they were again frozen overnight at −80 °C to further break up the tissue. In the morning, samples were centrifuged for 10 minutes at 14,000 x G. 10 μL of supernatant was added to a new transparent microcentrifuge tube followed by 50 μL of ProMega luciferin assay reagent immediately prior to reading the luminescence on a ProMega GLOMAX 20/20 Luminometer. Luminescence was normalized to the mass of tissue (luminescence/g tissue).

## Results

In this work, we based our formulation composition and components on the established Moderna LNP vaccine formulation which was used in the delivery of COVID-19 mRNA vaccines (**Supplemental Table S1**). To replace the immunogenic DMG-PEG lipid component, we evaluated a series of new polymers based on PMeOx and PEtOx with varying degrees of polymerization which were terminated with sodium azide to enable direct reaction with DBCO functionalized 16:0 PE phospholipids. The polymers ranged from targeted degrees of polymerization of 10-80 (**Table 1**), centered around the molecular mass of PEG (2000 g/mol) and the degree of polymerization (∼40) in the standard Moderna-like formulation **(Supplemental Table S2**). We also prepared a PEG-PE LNP formulation for direct comparison. In **Supplemental Figure S1A**, we demonstrate that during our LNP formulation process development steps, LNPs formulated with DMG-PEG and PE-PEG had comparable or even slightly superior *in vitro* and *in vivo* transfection levels when compared to DMG-PEG. Because of these initial findings during our optimization steps, we opted to continue the rest of this work with PE-PEG LNPs as our positive and clinically relevant control to better match the DBCO-phospholipid base in our POx LNPs. Additionally, we briefly varied the molar percentage of the PE-POx lipid with PMeOx-DP20 to see how it influenced transfection (**Supplemental Figure S1B**). As was previously shown pegylated LNPs^21,22^, increased molar percentage decreased the size but led to decreased *in vitro* transfection, so we opted to maintain a 1.5 mol % for the polymer-lipid throughout this work.

**Table 1:** Characterization of polymers synthesized for this study centered around the molecular weight and degree of polymerization of PEG-2000. Individual polymer batches were synthesized.

| Polymer | Batch Code | Nominal DP | HNMR DP | HNMR $M_n$ | GPC $M_n$ | GPC $M_w$ | PDI |
| --- | --- | --- | --- | --- | --- | --- | --- |
| PMeOx | V6 | 10 | 11 | 992 | 673 | 678 | 1.007 |
|  | V4 | 20 | 18 | 1587 | 1004 | 1083 | 1.039 |
|  | V5 | 40 | 56 | 4817 | 2832 | 3929 | 1.167 |
|  | V7 | 80 | 85 | 7282 | 4976 | 7633 | 1.275 |
| PEtOx | V3 | 10 | 11 | 1146 | 972 | 995 | 1.024 |
|  | V1 | 20 | 24 | 2433 | 1887 | 2003 | 1.061 |
|  | V2 | 40 | 57 | 5700 | 5686 | 6125 | 1.077 |
|  | V4 | 80 | 96 | 9561 | 13562 | 14609 | 1.077 |

### Polymer Synthesis

POx polymers were synthesized by living cationic ring opening polymerization (LCROP) which produces well-defined, low dispersity polymers. **Table 1** shows the polymers which were synthesized including their nominal DP values, actual DP values, and molecular masses as determined by ^1^HNMR and GPC along with polymer PDI. Confirmation of azide group on purified polymers was assessed by FTIR. **Figure 1** shows representative ^1^HNMR, GPC, and FTIR characterization for the PEtOx-DP20 polymer. ^1^HNMR, GPC, and FTIR characterization for all synthesized polymers are found in **Supplemental Figures S2-S9.**

**Figure 1:**
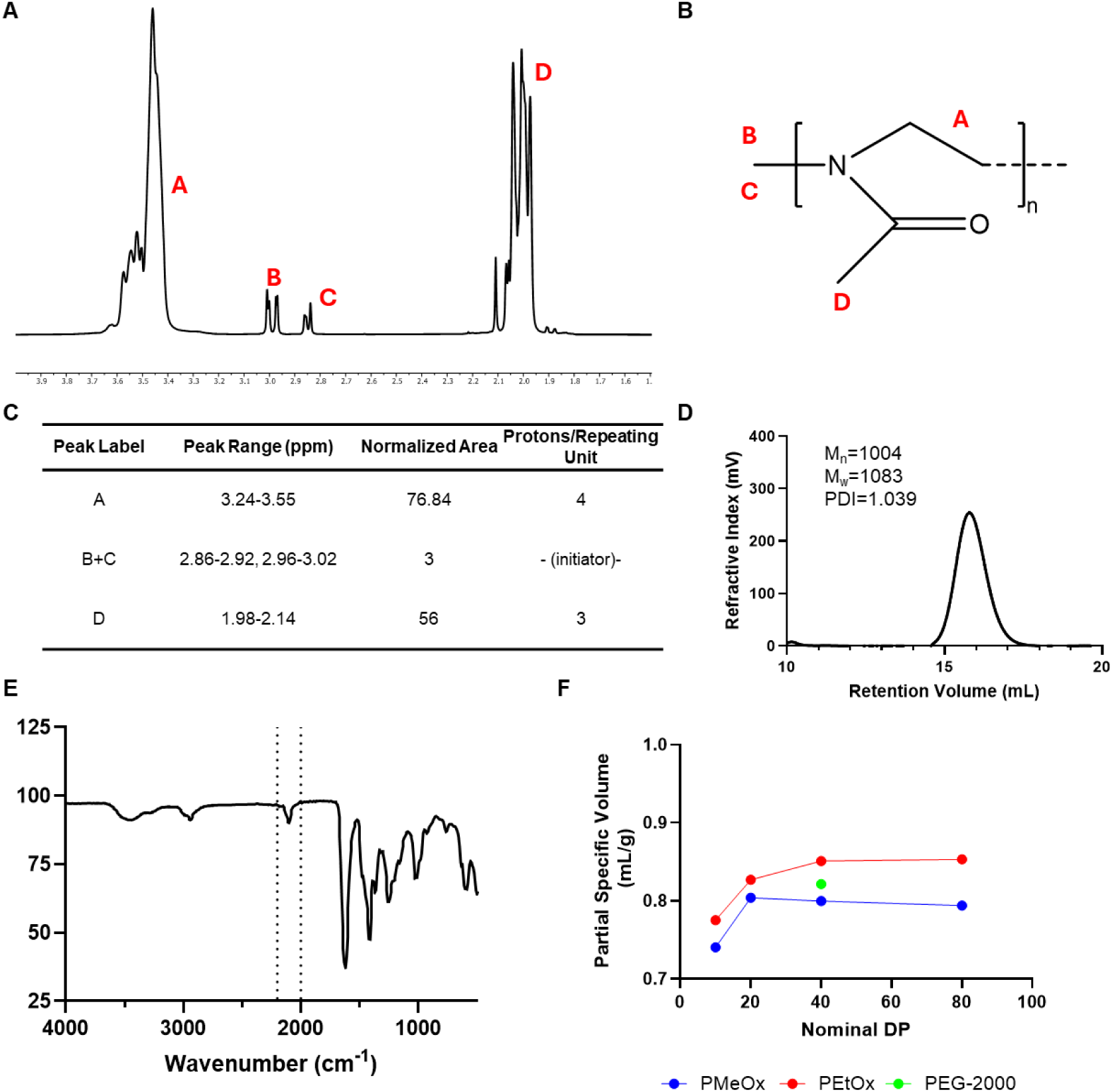
Characterization of the nominal DP-20 PMeOx polymer representative of characterization for all polymers synthesized in this work b (A) ^1^HNMR with peaks labeled corresponding to the (B) molecular structure. (C) Table of NMR peaks, their ranges on the spectrum, the normalized area, and number of protons per repeating unit used to estimate DP and M_n_ molecular weight. (D) GPC trace indicating M_n_, M_w_, and PDI. (E) FTIR spectrum of polymer indicating characteristic azide peak from termination with sodium azide at ∼2100 cm^-1^. (F) The partial specific volumes measured for each polymer investigated in this work.

### Polymer-Lipid Conjugation

Polymers and lipids were mixed in a 3:1 polymer:lipid molar ratio in ethanol to produce POx conjugated PE. The DBCO group on the PE has a characteristic UV absorbance peak around 310 nm. After reacting with the azide group at the end of the polymers, the peak disappears. This conjugation was performed with ethanol as a solvent, and polymer lipids were stored at –20 °C in ethanol long term. This signal disappears quite rapidly over 1-3 hours, but the reaction was allowed to proceed overnight before utilizing the lipids (**Supplemental Figure S10**). The absorbance at 310 nm is shown to decrease completely after overnight reaction in both PMeOx (**S10A**) and PEtOx (**S10B**) polymers.

### LNP Formulation

LNPs were formulated utilizing a Helix Bio impinged jet mixer. We prepared four batches of PEG LNPs and two individual batches for each POx polymer studied in this work to assess inter-batch variability and reproducibility of key quality attributes for LNPs. We used DLS to assess intensity and number average size as well as particle PDI. All DLS histograms for individual batches are presented in **Supplemental Figure S11**. Altogether we produced 20 batches of LNPs and their characteristics are summarized in **Supplemental Table S3**. For the most part, all batches produced similar numbers of particles as measured by NTA. The particle sizes were generally consistent across formulations with low PDI. Notably, the LNPs prepared using the shortest DP10 polymers, both PMeOx and PEtOx, exhibited larger diameters showing an increase of approximately 10–15 nm in particle size. This is consistent with previous observations on pegylated LNPs of various lengths^23^. We noted that as the POx degree of polymerization increased, we saw improved recovery of LNPs (as measured by the NTA particle counts, **Table 2**), which indicates that steric hindrance from extended polymer chains may reduce unwanted material interactions during LNP preparation.

**Table 2:** Summary data for all LNP formulations prepared (Mean±SEM presented). Four batches of PEG LNPs were prepared and two batches of each POx LNP were prepared to assess inter-batch variability.

| LNP Formulation |  | Dynamic Light Scattering (DLS) |  |  | Nanoparticle Tracking Analysis (NTA, Zetaview) |  |  |  |  | RiboGreen RNA Quantification Assay |  |
| --- | --- | --- | --- | --- | --- | --- | --- | --- | --- | --- | --- |
| Polymer | Nominal DP | Intensity Size | Z-Average Size | PDI | Mean Size | Median Size | Mode Size | Total Particles | Zeta Potential | Total mRNA Concentration | Encapsulation Efficiency |
|  |  | (nm) | (nm) | - | (nm) | (nm) | (nm) | - | (mV) | (ug/mL) | - |
| PEG | MW 2000 g/mol | 98.0±5.84 | 88.6±5.85 | 0.122 ± 0.014 | 77.5±2.179 | 77.5±1.5 | 75.8±1.3 | 1.0E+12±1.6E+11 | -5.45 ±0.60 | 252.0±10.4 | 91.9%±0.6% |
| PMeOx | DP 10 | 135.8±6.1 | 121.3±1.1 | 0.180 ± 0.008 | 87.5±0.200 | 87.5±0.6 | 82.5±0.5 | 7.4E+11±1.9E+10 | -9.42 ±1.48 | 203.5±16.6 | 85.1%±1.5% |
|  | DP 20 | 100.8±4.7 | 91.4±2.7 | 0.178 ± 0.004 | 74.8±1.900 | 74.8±1.3 | 72.5±0.8 | 9.5E+11±3.6E+11 | -4.90 ±0.79 | 212.9±5.8 | 88.0%±1.0% |
|  | DP 40 | 81.2±0.9 | 74.6±0.2 | 0.083 ± 0.018 | 76.0±0.300 | 76.0±0.8 | 74.8±0.7 | 1.1E+12±1.1E+09 | -3.06 ±0.06 | 205.2±26.2 | 85.8%±0.4% |
|  | DP 80 | 87.9±1.2 | 80.1±0.2 | 0.085 ± 0.014 | 77.6±0.350 | 77.6±0.6 | 76.5±0.6 | 1.1E+12±4.2E+10 | -3.79 ±1.06 | 184.4±1.1 | 85.0%±3.9% |
| PEtOx | DP 10 | 129.8±24.8 | 112.5±17.5 | 0.172 ± 0.011 | 83.8±8.350 | 83.8±6.0 | 79.0±3.4 | 8.6E+11±2.5E+10 | -9.90 ±2.66 | 188.0±4.7 | 77.7%±0.0% |
|  | DP 20 | 89.8±2.3 | 82.7±0.8 | 0.182 ± 0.000 | 72.1±0.050 | 72.1±0.5 | 70.9±0.8 | 9.8E+11±2.4E+10 | -3.75 ±0.39 | 259.6±13.5 | 84.0%±0.0% |
|  | DP 40 | 83.4±2.9 | 74.2±0.9 | 0.142 ± 0.012 | 74.3±0.450 | 74.3±0.2 | 73.1±0.4 | 8.5E+11±3.7E+10 | -4.30 ±0.48 | 235.9±0.3 | 80.2%±1.3% |
|  | DP 80 | 92.4±6.1 | 82.8±4.0 | 0.189 ± 0.021 | 79.3±1.600 | 79.3±0.8 | 78.2±0.2 | 1.6E+12±8.7E+11 | -3.33 ±2.46 | 203.3±34.7 | 77.0%±2.3% |

All zeta potentials were slightly negative, and the incorporation of these neutral POx polymers had no discernible effect on zeta potential, regardless of polymer type or length. Similarly, the mRNA encapsulation efficiencies ranged from 77% to 91.9% with no clear trend observed as a function of polymer DP or chemical composition. Overall, our established manufacturing protocols consistently produced well-defined LNPs using the novel POx lipids, with very low inter-batch variability.

### *In Vitro* Transfection

To assess relative expression levels and potential cell-type tropism, we evaluated the in vitro transfection efficiency of our POx- and PEG-based LNPs across three cell lines: HEK293, DC2.4 (a murine dendritic cell line), and 4T1 (a murine triple-negative breast cancer cell line). Each individual batch was independently tested in these cell lines to evaluate any batch-to-batch variation in transfection efficiency. **Supplemental Figure S12** shows that overall, transfection levels were mostly consistent across batches, though some statistically significant differences were observed, particularly with the PEG formulations

Figure 2 **A, B, C,** and **D** shows the average transfection efficiency of the batches for each LNP formulation. Across all three cell types, the DP10 PMeOx and DP10 PEtOx formulated LNPs show consistently high transfection efficiency, comparable or superior to PEG-2000 LNPs in each cell line. The DP20 polymers generally show strong transfection as well but yielded statistically lower transfection than their DP10 counterparts in all three cell lines. As the DP of the polymers increases to 40 and beyond, transfection efficiency declines sharply. Across the tested cell lines and selected polymers, we observe a consistent trend of decreasing transfection levels from HEK293 to DC2.4 to 4T1, suggesting cell-type-dependent differences in uptake or expression efficiency.

**Figure 2:**
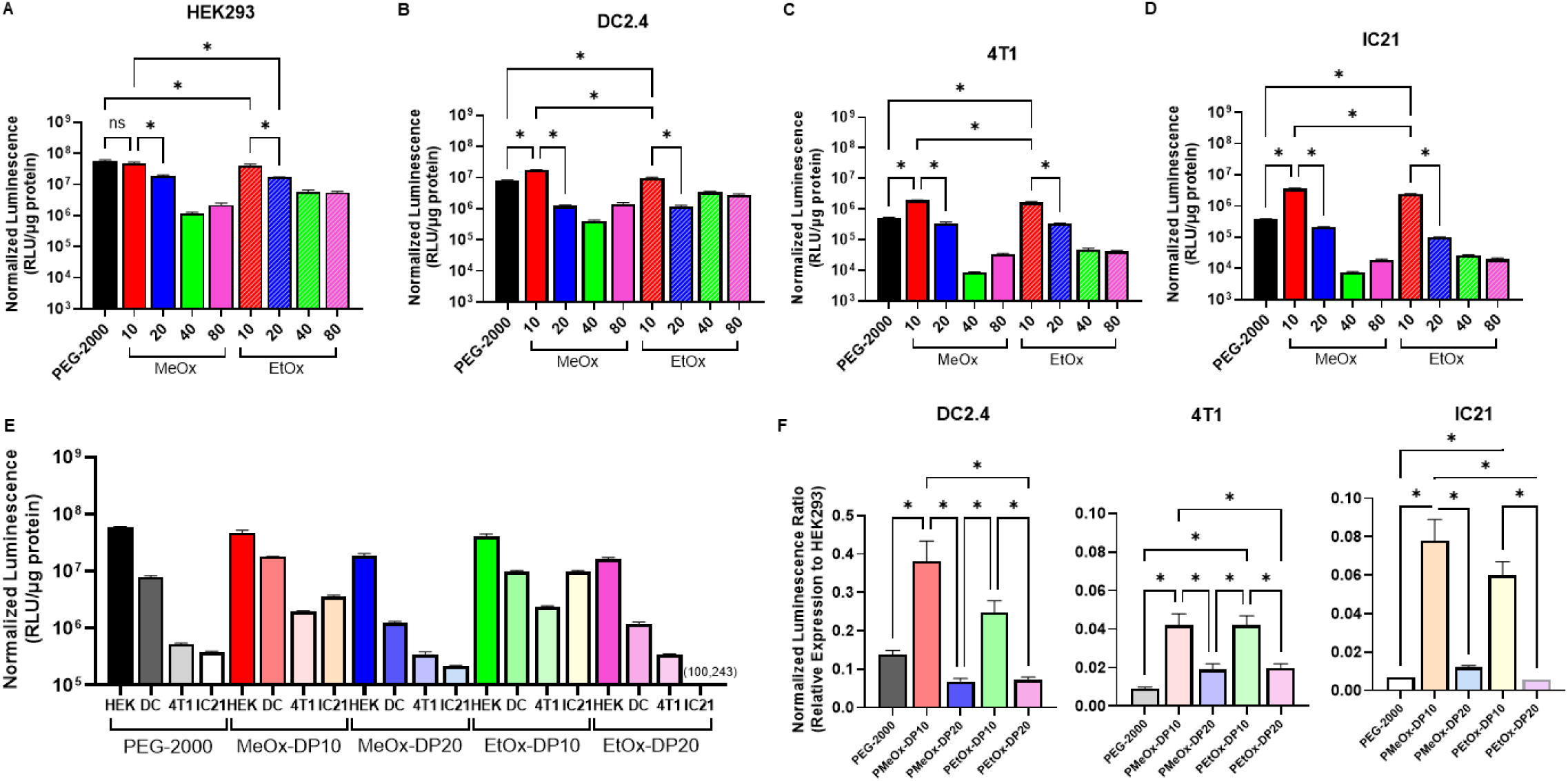
*In vitro* transfection of multiple cell lines including HEK293, DC 2.4, 4T1, and IC21 **(A)** Normalized luminescence from firefly luciferase mRNA transfection after 24 hours in HEK293 cells (n=4 PEG-2000 batches, n=2 Oxazoline batches). **(B)** Normalized luminescence from firefly luciferase mRNA transfection after 24 hours in DC 2.4 cells (n=4 PEG-2000 batches, n=2 Oxazoline batches). **(C)** Normalized luminescence from firefly luciferase mRNA transfection after 24 hours in 4T1 cells (n=4 PEG-2000 batches, n=2 Oxazoline batches). **(D)** Normalized luminescence from firefly luciferase mRNA transfection after 24 hours in IC21 cells (n=4 PEG-2000 batches, n=2 Oxazoline batches). **(E)** Comparison of select polymers with consistently high expression levels *in vitro* (PEG-2000, MeOx-DP10, MeOx-DP20, EtOx-DP10, and EtOx-DP20) in all four cell lines. **(F)** The ratio of DC 2.4, 4T1, and IC21 transfection levels to HEK293 expression with select polymers with consistently high expression levels *in vitro* (PEG-2000, MeOx-DP10, MeOx-DP20, EtOx-DP10, and EtOx-DP20) indicating some polymer, and polymer length-dependent LNP tropism in each cell type cell transfection.

Within each LNP formulation type, all cell types very different levels of transfection (Figure 2E). To further investigate potential differences in cell-type tropism, we compared transfection profiles across formulations, aiming to identify any preferential targeting or expression patterns among HEK293, DC2.4, 4T1, and IC21 cells. All LNP formulations were relatively poor at transfecting 4T1 an IC21 cells compared to HEK293 and DC2.4 cells (luminescence ∼1% or less of HEK293 in many groups). Using HEK293 as a baseline expression level, where we expect no formulation to have inherent preferential tropism, we compared the ratio of DC2.4, 4T1, and IC21 transfection to HEK293 (Figure 2F). We observed that the short PMeOx DP10 polymer exhibited increased DC2.4, 4T1, and IC21 to HEK293 transfection ratios compared to PEG-based LNPs (one-way ANOVA analysis, α=0.05). The DP10 PEtOx polymer showed significantly improved transfection ratios as well in the 4T1 and IC21 cell lines (one-way ANOVA analysis, α=0.05). The DP20 PMeOx and PEtOx polymers exhibited not significantly different ratios in each of the cell lines. Generally, the short DP10 polymers outperformed the DP20 polymers of the same kind. This data suggests an enhanced propensity of these short-chain POx LNPs to transfect dendritic cells, cancer cells, and macrophages relative to other formulations, like the pegylated LNPs. These improvements of short POx lipids to transfect dendritic cells highlights their possible utility and advantages over the standard PEG-LNP in a vaccine platform.

### *In Vivo* Biodistribution and Transfection

While *in vitro* and *in vivo* transfection efficiencies can be correlated, this is not always the case. All tissue types face drug delivery barriers and may transfect differently compared to the *in vitro* systems. To this end, we wanted to evaluate our ability to express mRNA in a vaccine setting. Formulations for injection were prepared by mixing the two individually formulated batches into a single solution for each formulation. We injected balb/c mice with 5 μg of mRNA loaded in LNPs intramuscularly in the right gluteus medius muscle. We selected the top performing LNPs *in vitro* for these experiments including DP 10 and DP 20 PMeOx and PEtOx LNPs. We also selected DP40 PEtOx and used the PEG-2000 LNPs as a clinically relevant control. Figure 3 shows the *in vivo* luminescence images for each of the formulations. Overall, expression is high across all formulations, with PMeOx and PEtOx DP10 having some localized hot spots of increased transfection. Additionally, from examining the images in Figure 3A, there appears to be a greater tendency for expression to spread outside of the muscle, into tissues other than just the liver, for the DP10 polymers compared to PEG-2000 and DP20 polymers. PEG-2000 and DP20 polymers appear to have luminescence more strictly confined to muscle and liver tissues. Figure 3A shows the luminescence from a side-view of each mouse while Figure 3B shows a ventral view where we could appropriately monitor full-body and liver luminescence. Notably, for the PEG-2000 based LNPs in Figure 3B the expression in the liver increases from 4 to 24 hours post injection. Meanwhile, for each of the POx-based LNPs, the liver expression decreases from 4 to 24 hours post injection, except the short DP10 PMeOx polymer in males, which stayed relatively stable.

**Figure 3:**
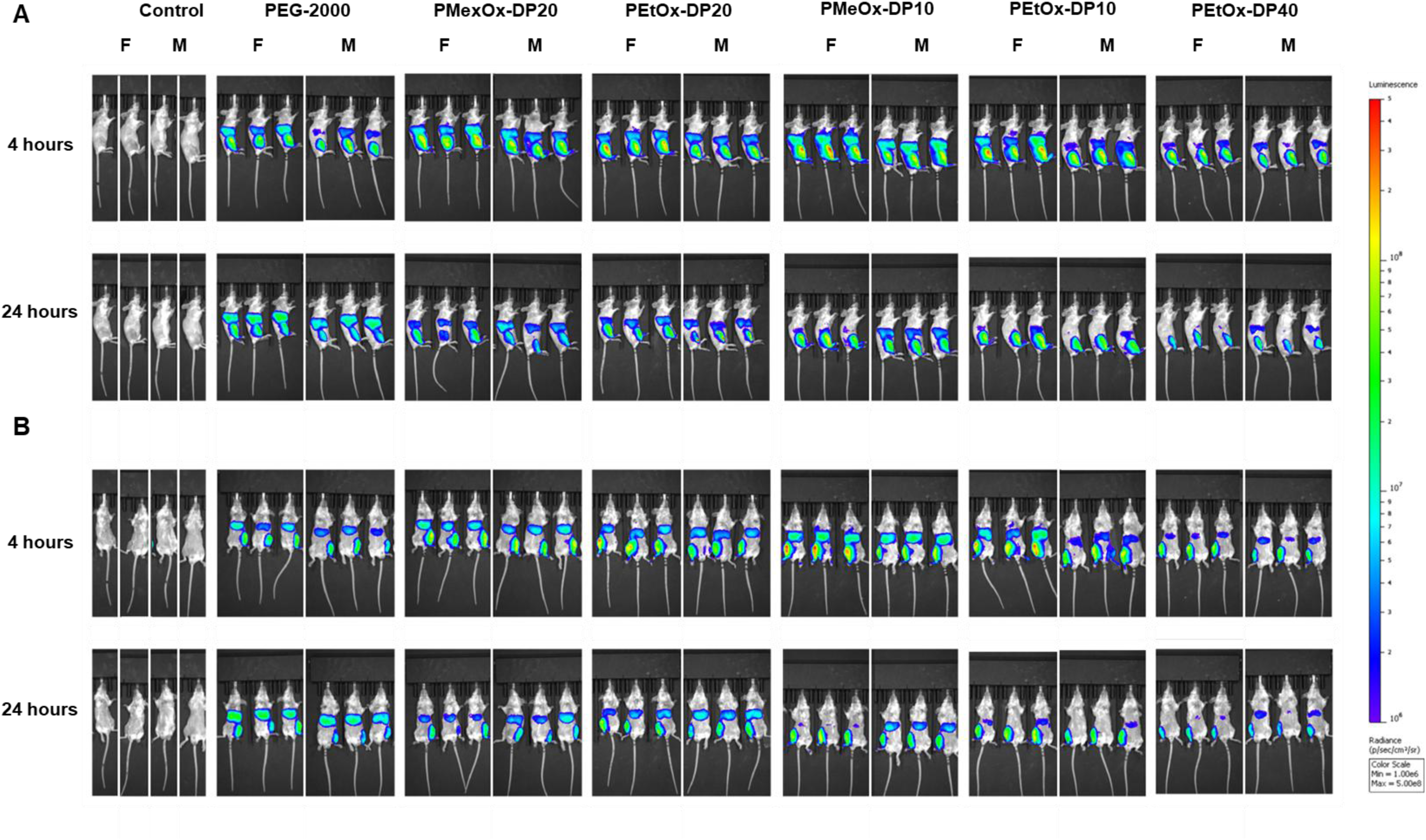
*In Vivo* luminescence analysis of full body luciferase expression in two views (side and ventral). **(A)** Side view showing expression in right gluteus medius muscle (and liver) at 4 and 24 hours post injection where LNPs were injected. Muscle luminescence quantification was done using these images. **(B)** Ventral view showing expression in the liver (and right gluteus medius muscle) at 4 and 24 hours post injection with LNPs. Liver luminescence quantification was done using these images.

**Supplemental** Figure 13 shows sex-based differences in luminescence across tissues and time points. Muscle expression is shown in the side view images (Figure 3A), while liver expression is clearly viewed in the ventral view images (Figure 3B). Statistically significant intersex differences were observed with several of the LNP formulations in both the muscle and liver at each time point (one-way ANOVA with selected comparisons between sexes within each formulation, α=0.05). Beyond statistical significance, a consistent trend emerged: in most formulations, muscle expression levels were higher in female mice compared to male mice. Male mice showed a trend of higher liver accumulation compared to females.

This dataset was further analyzed in Figure 4, where muscle and liver expression levels were compared across treatment groups within each sex to evaluate how the novel POx LNPs compare to the standard PEG-2000 LNPs. Figures 4A and **4B** present muscle expression in female and male mice, respectively, at 4 and 24 hours post injection. Figures 4C and **4D** show the corresponding liver expression data. At the 4-hour time point, both PEtOx and PMeOx polymers with DP-10 show increased muscle expression in female mice while only PEtOx DP10 showed a similar enhancement in male mice. By 24 hours, both polymers improved muscle expression in males, suggesting a time-dependent effect. In contrast, liver expression in PEG-2000 LNP-treated mice increased from 4 to 24 hours in both sexes. Notably, POx LNPs generally exhibited a decrease in liver expression over the same time period, indicating a potentially favorable biodistribution profile for vaccine applications.

**Figure 4:**
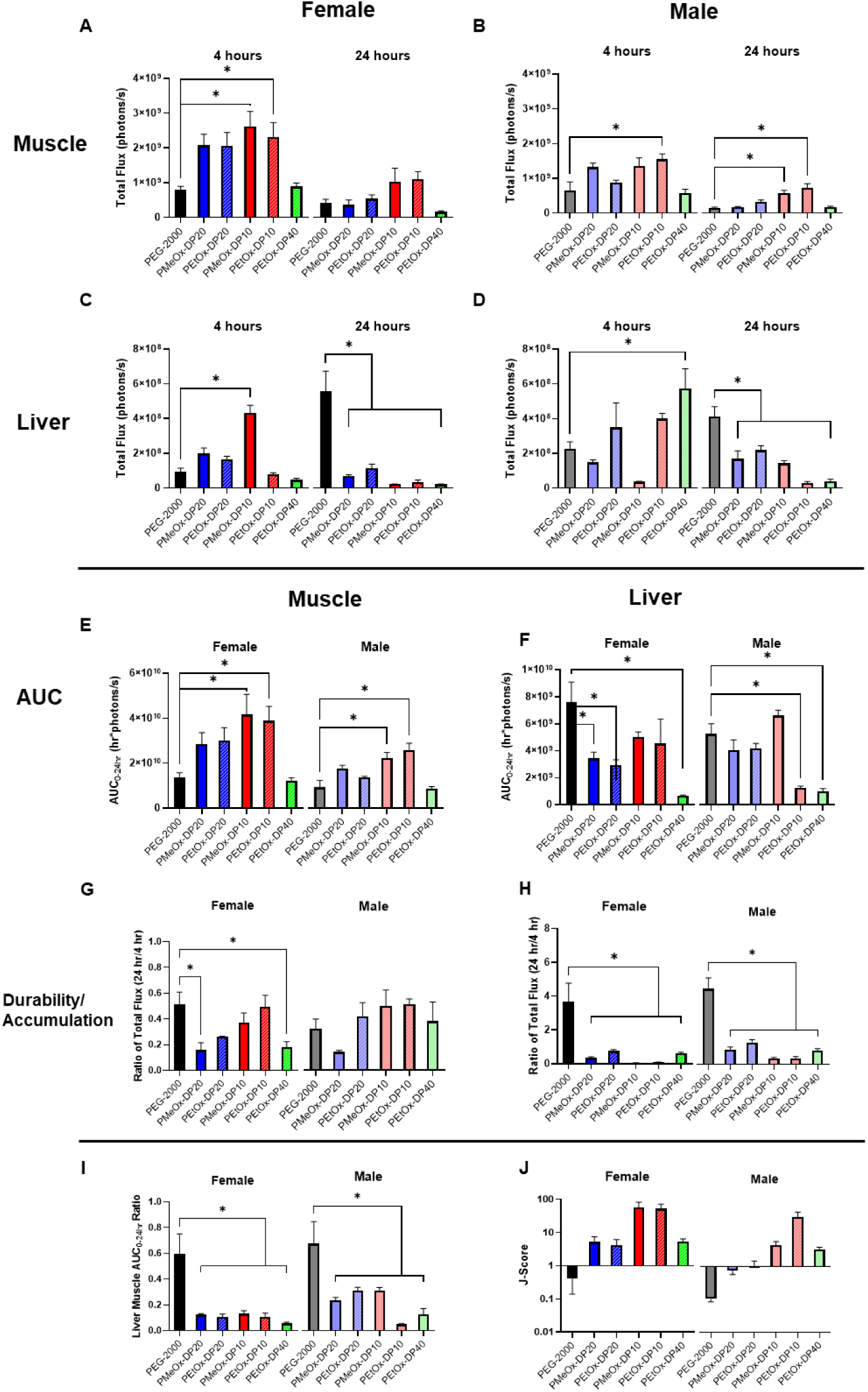
Quantification of *in vivo* IVIS full body imaging analyzing muscle and liver tissue expression. Female and Male were analyzed separately and individual metrics analyzed by one way ANOVA comparing all columns to the PEG-2000 standard formulation (α=0.05, N=3 for per treatment/gender). **(A,B)** Female and Male muscle expression at 4 and 24 hours **(C,D)** Female and Male Liver expression at 4 and 24 hours. **(E,F)** Female and Male AUC from 0-24 hr in the muscle and liver. **(G)** Muscle transfection durability in female and male mice (ratio of 24 hr expression to 4 hour expression) **(H)** Liver transfection accumulation in female and male mice (ratio of 24 hr expression to 4 hour expression) **(I)** Ratio of Liver AUC to Muscle AUC in female and male mice. **(J)** J-Score metric for comparing formulations.

Figure 4E shows that the muscle area under the curve (AUC) for both DP10 POx polymers was significantly higher than for PEG-2000 LNPs. Additionally, DP20 POx polymers demonstrated reduced liver accumulation in female mice (Figure 4F). Although PMeOx-DP10 LNPs exhibited initially high liver uptake, this decreased drastically by 24 hours post-injection. To further assess biodistribution, we calculated the AUC ratio (AUC-Liver/AUC-Muscle) as a measure of each formulation’s tendency to avoid liver accumulation—a factor known to reduce vaccine efficacy and increase toxicity (Figure 4 **I**)^7^.

While several statistically significant improvements were observed POx LNPs compared to PEG-2000 LNPs, we sought to integrate these findings into a single composite metric that captures key performance attributes: sustained expression in muscle tissue, reduced liver accumulation, and overall expression levels. To this end, we developed the “J-Score”, a quantitative index that incorporates muscle durability (defined as the ratio of muscle gene expression at 24 hours to that at 4 hours; Figure 4G) and liver accumulation (defined as the ratio of liver gene expression at 24 hours to 4 hours; Figure 4H), as outlined in **Equation 2**.

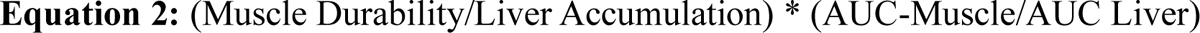

A higher J-Score indicates superior LNP performance, reflecting both sustained expression in the target tissue and reduced off-target accumulation. This metric enables direct comparison of formulation biodistribution by integrating multiple biodistribution and expression parameters into a single, interpretable value. As shown in Figure 4J, LNPs formulated with PEtOx-DP10 polymers achieved the highest J-Scores in both female and male mice, highlighting their strong expression profiles and reduced liver exposure—key attributes for effective and safe vaccine delivery. The J-Score can be further modified to incorporate overall expression levels for formulations with drastically different expression levels (**Equation 3**).

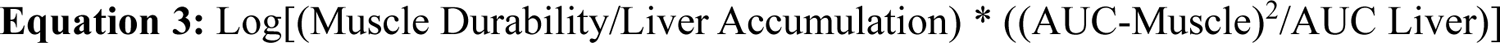

When comparing the J-Score to the Modified J-Score (**Supplemental Figure S14**), the conclusions are the same, because the expression levels, as determined by the live full body imaging, are on the same order of magnitude. For evaluated formulations which have expression levels varying by an order of magnitude, it is essential to utilized the modified J-Score. Because the J-Score only uses AUC ratios and Durability/Accumulation. It is somewhat agnostic to the absolute expression intensity. By using the modified J-Score, a score difference of 1-unit would occur if AUC ratios, muscle durability, and accumulation were the same but absolute muscle intensity was one order of magnitude less. In this work, we try to optimize muscle intensity and mitigate the liver accumulation. But this J-Score can easily be adapted to any two organs of interest to evaluate formulation differences in a single metric.

At 24 hours post-injection, mice were euthanized, and major organs were harvested for *ex vivo* luminescence analysis. **Supplemental** Figure 15 presents normalized luminescence values (RLU/g tissue) for both female and male mice, allowing visualization of expression across organs. To further explore trends in organ-specific expression as a function of LNP formulation, Figure 5A-H display individual organ data for both sexes, enabling direct comparison and identification of any sex-dependent differences in biodistribution.

**Figure 5:**
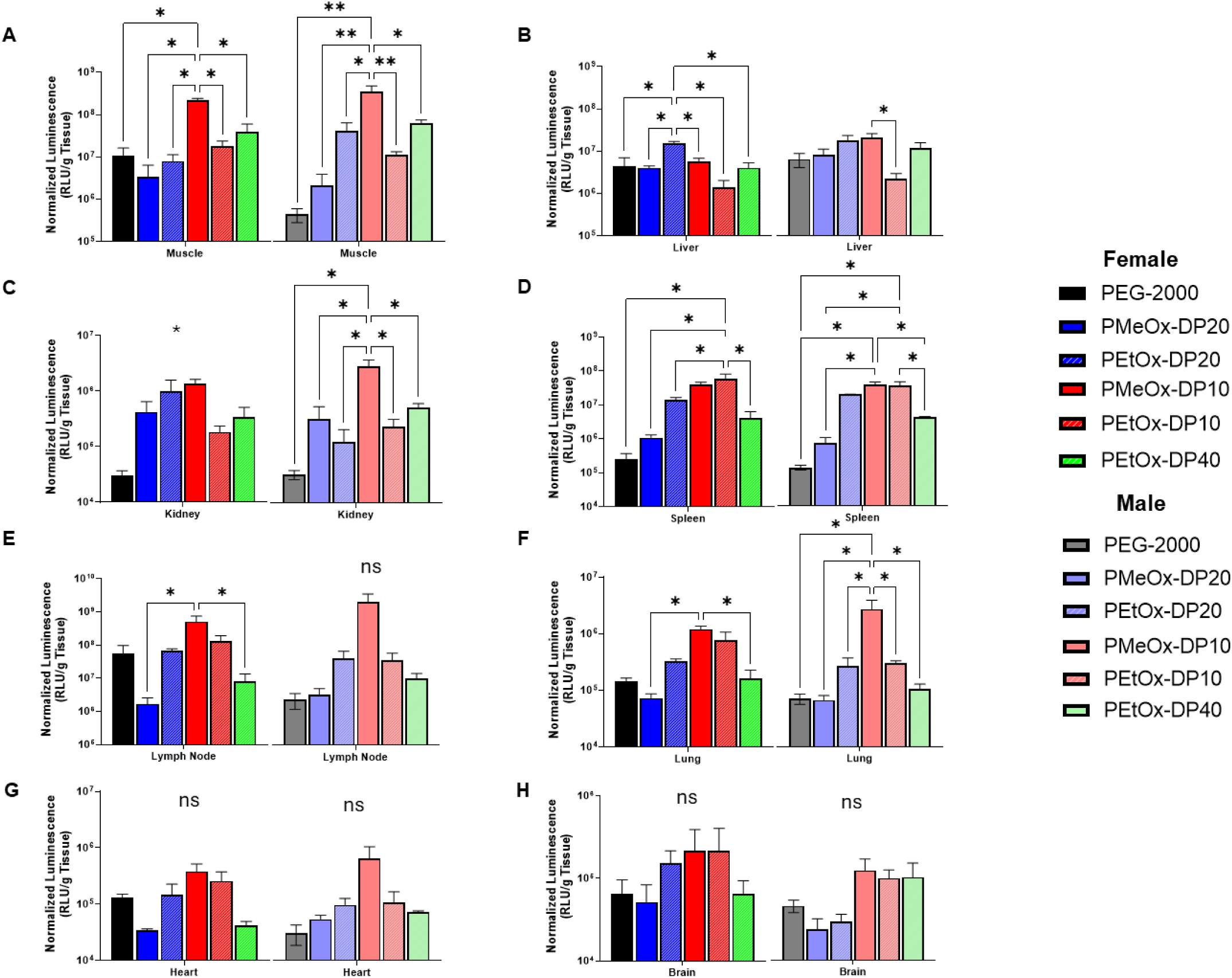
Normalized *ex vivo* luminescence (RLU/g Tissue) in excised organs at 24 hours post intramuscular injection as a function of LNP formulation and separated by gender **(A)** Muscle **(B)** Liver **(C)** Kidney **(D)** Spleen **(E)** Lymph Node **(F)** Lung **(G)** Heart **(H)** Brain. Statistical analysis was performed with a one-way ANOVA comparing all columns to each other within each tissue and gender (α=0.05). In female kidneys, ANOVA reports there is statistical significance between groups, but not pairwise comparisons.

In muscle expression, similar to the pattern observed with full-body luminescence, the shorter polymers appear to outperform the DP20 variants. In both females and males, the PMeOx-DP10 polymer shows significantly higher expression compared to all other polymers (Figure 5A). In the liver (Figure 5B), expression levels are relatively consistent across formulations, with no dramatic differences. However, in females, PEtOx-DP20 exhibits statistically higher liver expression than the other formulations. This trend is not observed in males, where only PMeOx-DP10 shows higher expression than PMeOx-DP20. Notably, liver expression values remain within the same order of magnitude across all formulations, whereas muscle expression spans multiple orders of magnitude. In the kidney (Figure 5C), PMeOx-DP10 demonstrates higher expression in males compared to all other groups, while no significant differences are observed among females.

One of the key findings in this study is the differential expression observed in the spleen and lymph nodes (Figure 5D and **5E**). These organs serve as critical training grounds for the immune system, and enhancing expression in these areas is likely to improve vaccine efficacy and therapeutic outcomes. In females, the shorter DP10 polymers again demonstrate superior transfection efficiency, with PEtOx-DP10 showing significantly higher expression than most other polymers. In the lymph nodes, PMeOx-DP10 outperforms some formulations, and although no statistically significant differences are observed in males, PMeOx-DP10 still exhibits the highest transfection efficiency.

Overall, LNPs with the shorter PMeOx and PEtOx DP10 polymers demonstrate more effective tissue permeation compared to their longer-chain counterparts. This observation aligns with the full-body luminescent imaging results, which revealed broader tissue distribution for the shorter polymers. As a result, these formulations achieve higher expression levels in a wider range of tissues than both PEG-2000 LNPs and POx lipids with longer polymer chains. Lastly, we assessed intersex differences in organ-specific expression across all formulations (**Supplemental Figure S16**), providing further insight into whether or not there is sex-based variation in transfection amongst the formulations. The data indicates that there are almost no intersex differences of gene expression in the different tissues. The short PMestOx-DP10 polymer had differences in the liver and lung, but the biological significance may be minimal.

## Discussion

In this work, we have clearly demonstrated that POx lipids may have multiple advantages over PEG-lipids in LNP formulation outside of any established immunogenicity issues. Vaccination strategies rely on obtaining expression in key cells of interest like antigen presenting dendritic cells. Traditionally, these can be difficult cells for LNPs to target^24^. However, our initial *in vitro* results indicate that our low DP POx polymers have some dendritic cell tropism, with an increased tendency to transfect these cells when compared to PEG LNPs. This is further supported by the data showing individual tissue accumulation *in vivo* after 24 hours post injection. In this experiment, there was improved transfection/expression in both lymph nodes and the spleen for the PMeOx-DP10 and PEtOx-DP10 LNPs, respectively. Full body imaging in Figure 4 shows that these smaller polymers allow the LNPs to spread much more broadly, unlike other polymers which appear to have two main sources of luminescence-the muscle tissue (injection site) and the liver. This alone highlights the utility of our new POx LNPs in the vaccine space.

Pairing this data with trends in liver accumulation based on full body imaging show that the short PMeOx polymers were able to mitigate extensive liver accumulation while also increasing the expression levels in the muscle. In fact, PEG LNPs continued to accumulate in the liver up to 24 hours, as indicated by an increased signal between 4 and 24 hours. In contrast, all POx LNPs showed significantly reduced expression levels at 24 hours compared to their levels at 4 hours. Notably, the short DP10 polymers did have high initial liver readings, which may be do their improved propensity to transfect macrophages (Figure 2) relative to the other polymers. Altogether, this data suggests that a much lower portion of the dose may make it to the liver, mitigating liver toxicity concerns associated with current LNPs^5,7^. This indicates that POx LNPs could be promising outside of vaccines as well, particularly in spaces where repeat dosing regimens are required, and liver accumulation could be concerning. To compare these formulations, which had muscle expression levels within the same order of magnitude, we developed the “J-Score” metric, which evaluates a formulations ability to create durable, lasting muscle expression while mitigating liver accumulation.

These findings highlight a potentially critical parameter for selecting optimal formulations and guiding future decision-making. Notably, the two formulations favored by the J-Score, the DP 10 PMeOx and PEtOx, consistently demonstrated strong performance across multiple tissues. Both showed particularly promising expression in muscle, spleen, and lymph nodes. These trends were largely consistent across both female and male mice. Additionally, the DP10 PMeOx LNPs exhibited enhanced accumulation in the lungs compared to other formulations, further supporting their potential as lead candidates for systemic delivery applications.

Another phenomenon we observed early on in this work is the intersex differences in expression levels in the full-body imaging. While it was not always statistically significant, there was a clear trend that in many formulations, females exhibited 2-3X higher transfection levels compared to males. Additionally, the male mice tended to have higher liver accumulation compared to females. Overall thought, POx LNPs were able to reduce the liver/muscle AUC ratio over the observed 24-hour period and only two intersex differences in tissue expression were noted (**Supplemental Figure S16**). These intersex differences are a critical phenomena we need to be attentive to moving forward in this field to ensure transfection and vaccination efficacy is maintained across sexes as we explore novel polymer systems.

In mice, it is known that females have higher amounts of circulating immune cells and a different immune phenotype leading to a more efficient acute inflammatory response, which may be contributing to this phenomenon^25^. One prior study on sex differences in LNP vaccines in mice noted no difference in luciferase expression in the muscle, but significantly higher IgG responses in females compared to males^26^. This phenomena is partially recapitulated in humans, where the different sex hormones in females lead to differential innate immune responses and inflammasome reactions to different therapies and vaccines^27–29^. We postulate here that these differential immune populations and inflammatory responses could be leading to differential transfection seen in the full body imaging.

The tissue specific analysis in Figure 5 utilized harvested organs post-24 hours. There were no significant muscle differences in the 24-hour live body imaging, which is consistent with the tissue specific *ex vivo* analysis. Future work will evaluate intersex differences at earlier time-points when gene expression may be at its peak.

As a caveat to the full body imaging, it is primarily muscle cells which are transfected at the injection site^30,31^, so perhaps it is the larger male body weight/muscle size (∼21 g vs, ∼17 g females) in this study that led to more diffuse transfection and lower luminescence in the full body imaging. Although, it is postulated that mRNA expression in secondary lymphoid organs drives immune responses to LNP mRNA vaccines, highlighting the impact of our POx LNPs improving accumulation in the lymph nodes and spleen^31–33^.

The data for the muscle expression was consistent in both the *in vivo* whole-body imaging and the *ex vivo* organ imaging, with both indicating improved expression levels from some of the POx LNP formulations. However, the differences in liver exposure seen in the full body imaging were not as apparent in the *ex vivo* organs. This may be attributed to the many other tissues near the liver contributing to the observed signal in the *in vivo* full body imaging. Further studies are warranted to evaluate the extent of differences in the liver accumulation.

## Conclusions

This work presented a simple synthetic route for preparing POx lipids capable of reproducibly manufacturing LNPs. The LNPs stabilized with our short PMeOx and PEtOx polymers (∼DP10) showed widespread transfection at early timepoints, pervading many tissues and improving transfection/expression in the muscle, lymph nodes, and spleen. These LNPs showed an improved propensity transfect dendritic cells *in vitro* compared to standard PEG LNPs. Improved transfection efficacy and decreased liver accumulation makes these POx LNPs promising candidates for replacing PEG in LNP formulations moving forward. Altogether, this is a significant step towards improving the safety profile of LNPs in repeat administration settings and possibly improving vaccine efficacy by targeting key immune cell training grounds and antigen presenting cells.

## Acknowledgements

This work was supported in part by NIH T32CA196589 that provided fellowship to Dr. Colin Basham and 1U19 AI181979 consortium funding (AVK). Dr. Jacob Ramsey was supported by NCI F99CA274702 and K00CA274702. Additional funding from UNC Lineberger Center for Triple Negative Breast Cancer and Dr. Kabanov UNC startup funds. The authors would like to thank the Small Animal Imaging Core Facility, funded in part by the NCI Cancer Center Support Grant, P30CA016086, and the NIH S10 Shared Instrumentation Grant (NMR), S10OD026951, for supporting this work. The authors would like to acknowledge Dr. Son Long Ho for his insightful conversations on POx-lipid chemistry.

## Contributions

**Colin Basham:** Conceptualization, Data Curation, Methodology, Visualization, Writing, Review and Editing, formulation optimization, *in vitro* and *in vivo* studies. **Matt Haney:** Data Curation, Methodology, *in vitro* and *ex vivo* tissue data collection and analysis. **Yuling Zhao:** Data Curation, Methodology, *in vivo* data collection. **Konstantin A. Lukyanov:** Methodology, Review and Editing. **Kyoungtea Kim:** Methodology, *in vivo* studies. **Ayse Baysal:** Methodology, Polymer Characterization, *ex vivo* tissue analysis. **Hallie Hutsell:** Polymer Characterization, Review and Editing. **Alexander V. Kabanov:** Conceptualization, Writing, Review and Editing, Project Administration, Funding acquisition. **Jacob D. Ramsey:** Conceptualization, Data Curation, Methodology, Visualization, Formulation Characterization and Development, *in vivo* imaging, *ex vivo* tissues analysis, Writing, Review and Editing, Project Administration.

