## SupplementalData for "PEG-Free Tunable Poly(2-Oxazoline) Lipids Modulate LNP Biodistribution and Expression In Vivo after Intramuscular Administration"

**Supplemental Information**

**Supplemental Table S1:** Mole % lipids used in Moderna-like LNPs and in this work with lipid molecular weight (PEG-DMG listed in table)

| | Mol % | Molecular Weight (g/mol) |
|---|---|---|
| **Ionizable Lipid SM-102** | 50 | 710.2 |
| **Cholesterol** | 38.5 | 386.7 |
| **DSPC** | 10 | 790.2 |
| **PEG-Lipid (DMG for Moderna, 16:0 PE for this Work)** | 1.5 | 2509.2 |

**Supplemental Table S2:** PEG-2000 DP and molecular weight equivalent polymer characteristics for PMeOx and PEtOx

| | Molecular Weight (g/mol) | Degree of Polymerization (DP) |
|---|---|---|
| **PEG-2000** | 2000 | ~45.5 |
| **PMeOx (DP equivalent)** | ~3868 | 45.5 |
| **PMeOx (MW equivalent)** | 2000 | ~23.5 |
| **PEtOx (DP equivalent)** | ~4505 | 45.5 |
| **PEtOx (MW equivalent)** | 2000 | ~20.2 |

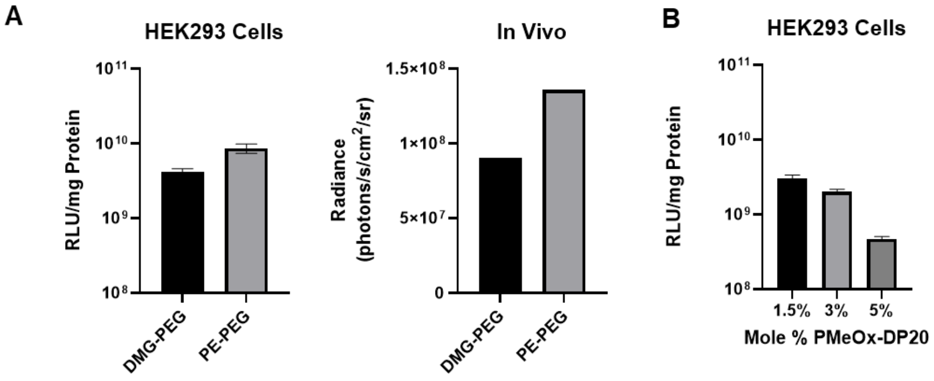


| Mol % POx Lipid | Size (d.nm) | PDI |
|---|---|---|
| 1.5 | 144 | 0.10 |
| 3.0 | 134 | 0.18 |
| 5.0 | 119 | 0.18 |

**Supplemental Figure S1:** Parameters evaluated in optimizing our LNP formulation **(A)** Comparison of DMG-PEG and PE-PEG in LNP formulation transfection *in vitro* in HEK293 cells and *in vivo* in single mice intramuscular injection and 24 hour luminescence imaging. **(B)** Evaluating the effect of varying the mol % polymer lipid with one of our lead PMeOx polymers (nominal DP20).

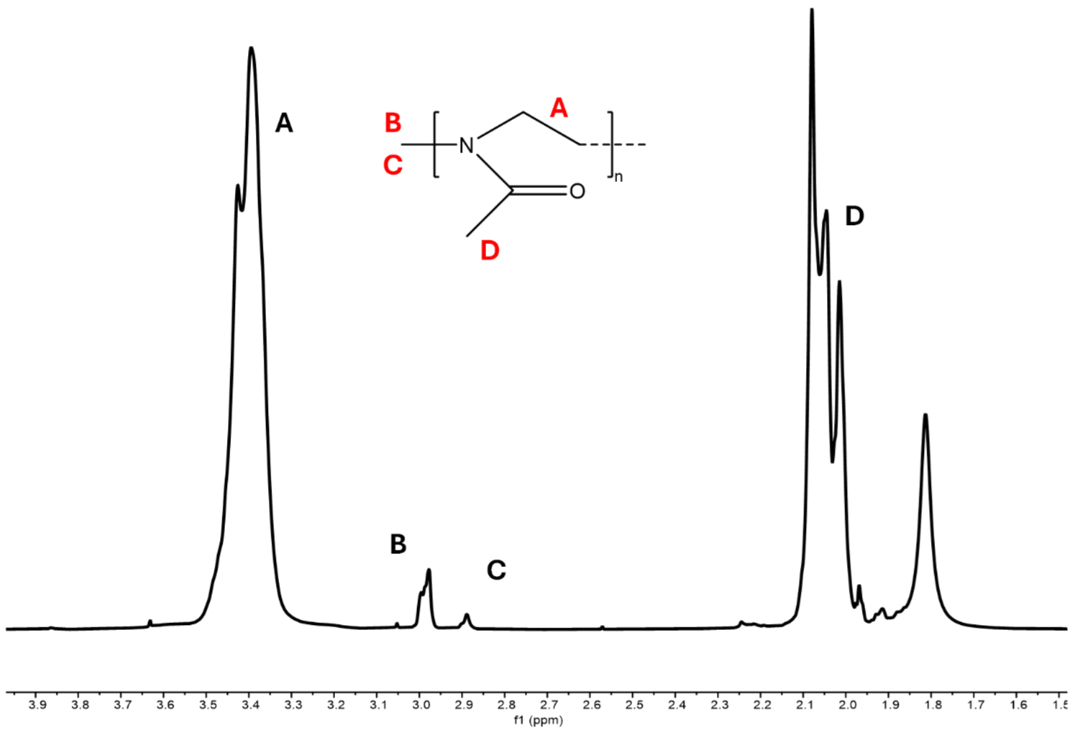


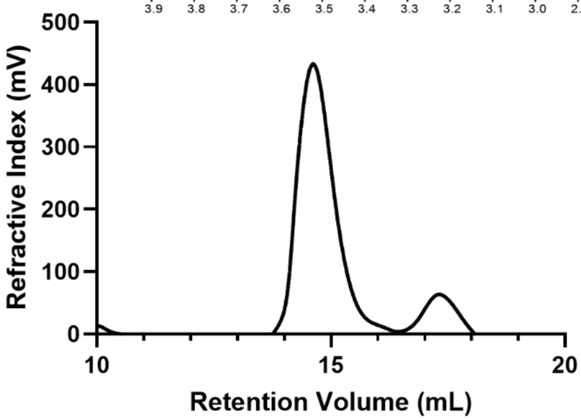


| Peak Label | Peak Range (ppm) | Normalized Area |
|---|---|---|
| A | 3.35-3.67 | 43.23 |
| B+C | 2.94-3.04, 2.81-2.89 | 3 |
| D | 1.94-2.09 | 30.95 |

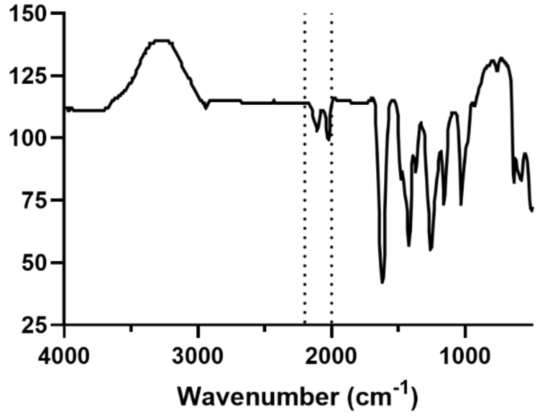


**Supplemental Figure 2:** $^{1}$HNMR, GPC, and FTIR Spectra indicating presence of azide peak from termination for PMeOx-V6 (Nominal DP-10).

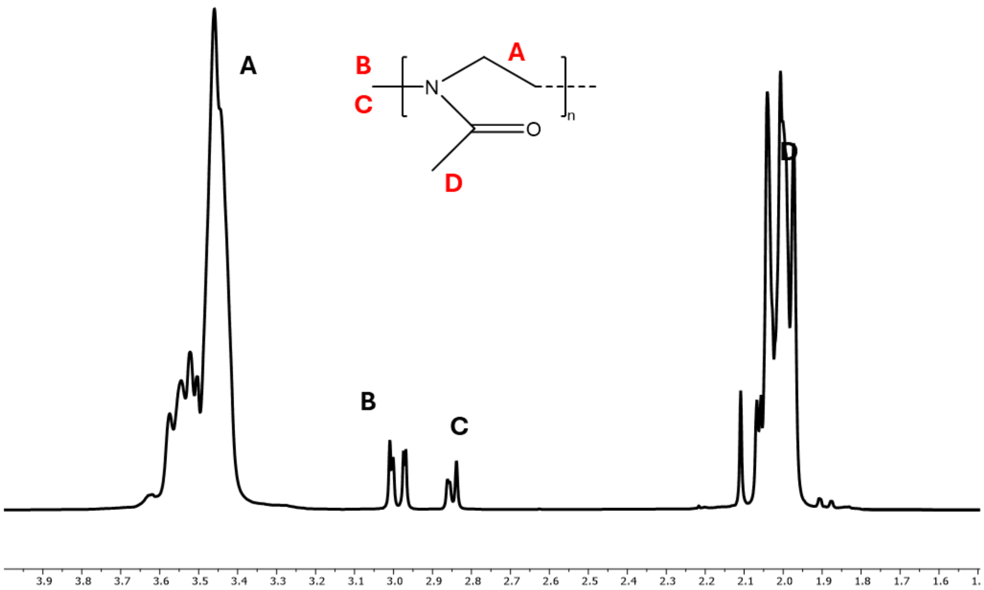


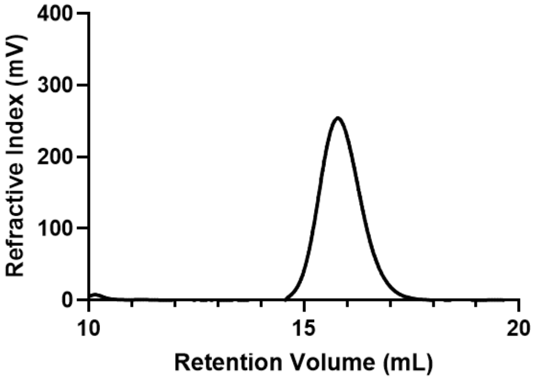


| Peak Label | Peak Range (ppm) | Normalized Area |
|---|---|---|
| A | 3.24-3.55 | 76.84 |
| B+C | 2.86-2.92, 2.96-3.02 | 3 |
| D | 1.98-2.14 | 56 |

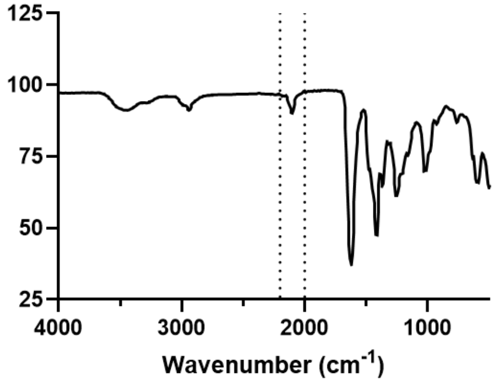


**Supplemental Figure S3:** $^{1}$HNMR, GPC, and FTIR Spectra indicating presence of azide peak from termination for PMeOx-V4 (Nominal DP-20).

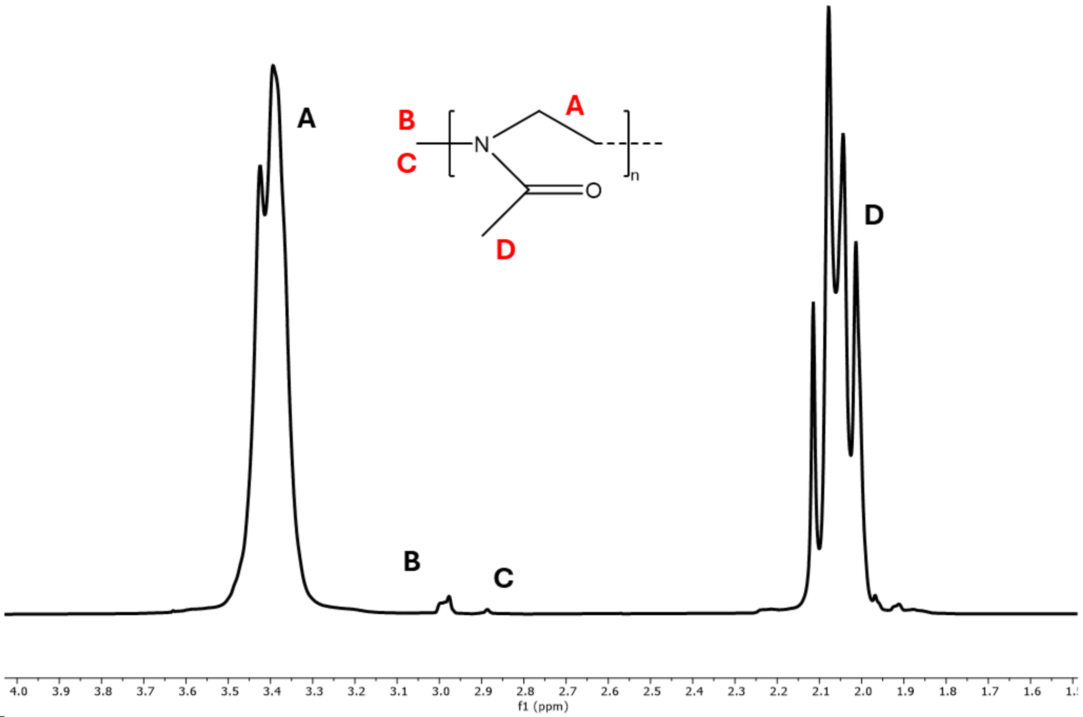


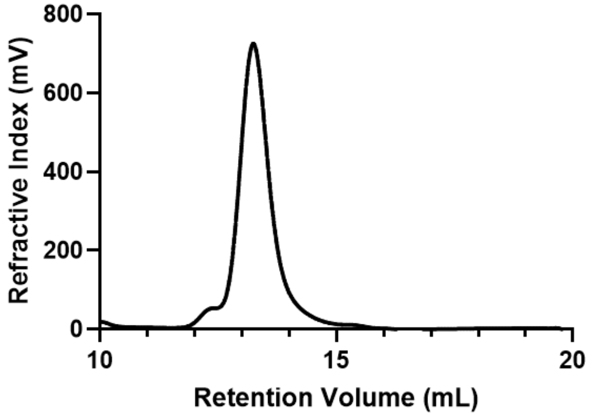


| Peak Label | Peak Range (ppm) | Normalized Area |
|---|---|---|
| A | 3.16-3.61 | 234.23 |
| B+C | 2.86-2.91, 2.96-3.02 | 3 |
| D | 1.98-2.10 | 168.77 |

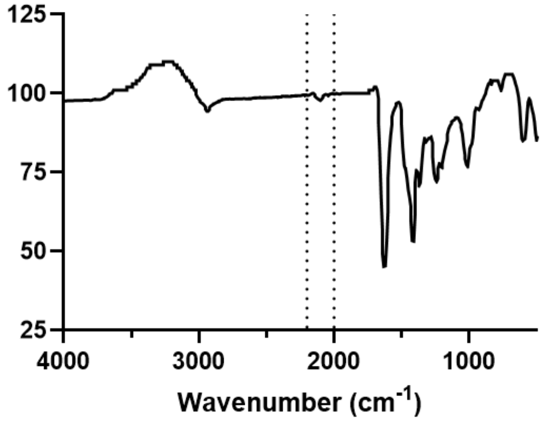


**Supplemental Figure S4:** $^{1}$HNMR, GPC, and FTIR Spectra indicating presence of azide peak from termination for PMeOx-V5 (Nominal DP-40).

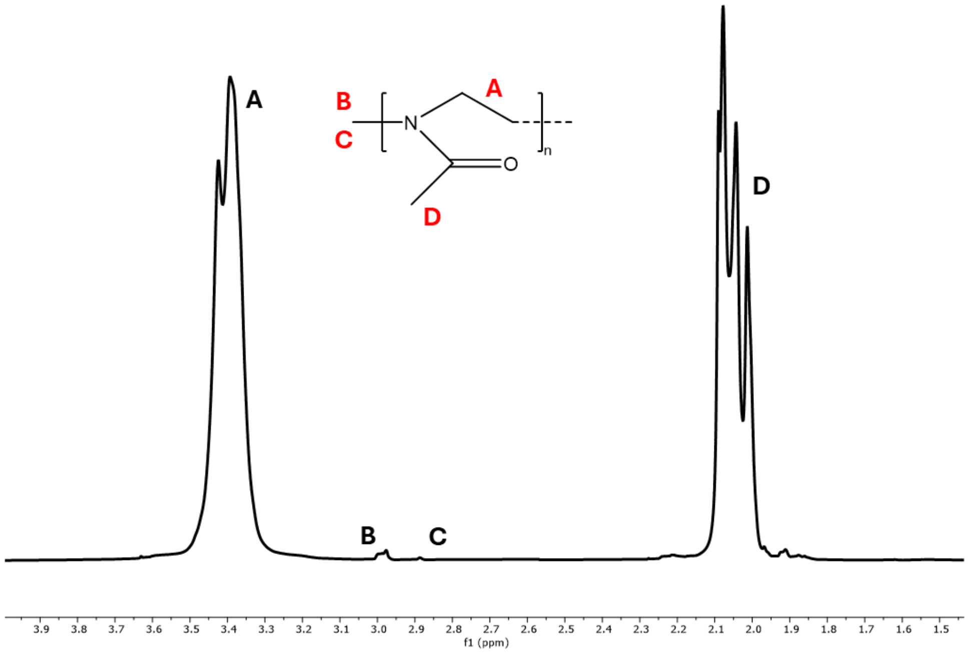


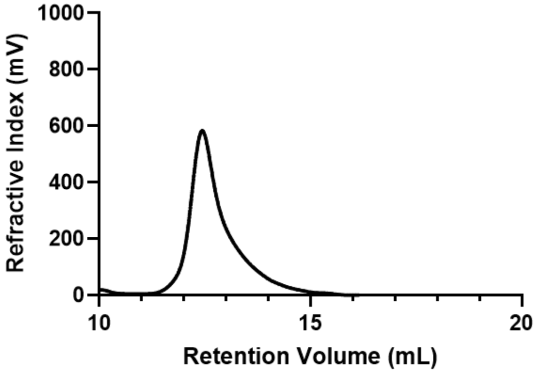


| Peak Label | Peak Range (ppm) | Normalized Area |
|---|---|---|
| A | 3.20-3.60 | 334.58 |
| B+C | 2.86-2.91, 2.95-3.03 | 3 |
| D | 1.98-2.13 | 272.50 |

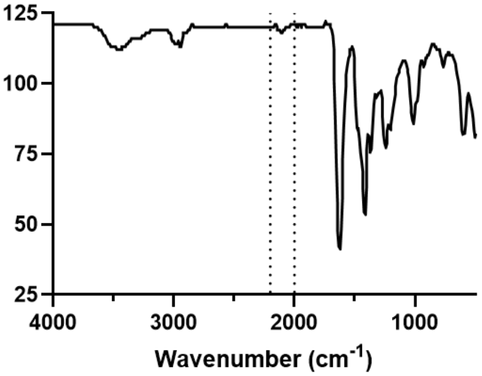


**Supplemental Figure S5:** $^{1}$HNMR, GPC, and FTIR Spectra indicating presence of azide peak from termination for PMeOx-V7 (Nominal DP-80).

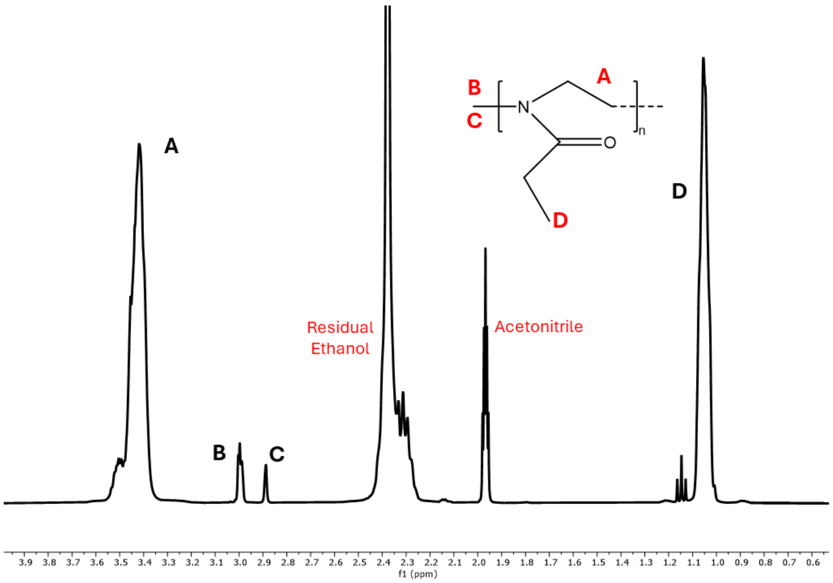


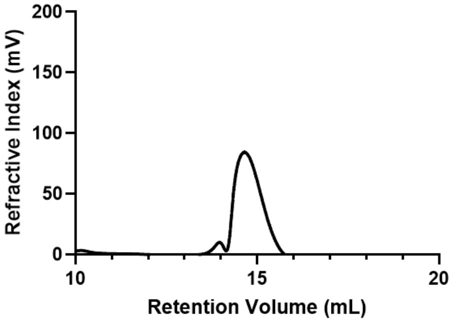


| Peak Label | Peak Range (ppm) | Normalized Area |
|---|---|---|
| A | 3.32-3.56 | 44.90 |
| B+C | 2.86-2.91, 2.97-3.02 | 3 |
| D | 0.99-111 | 33.28 |

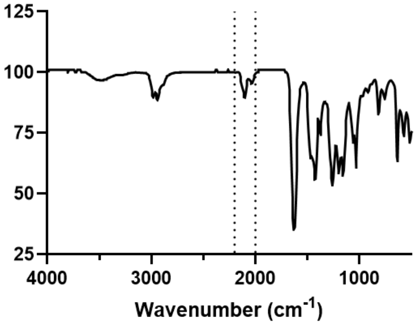


**Supplemental Figure 6:** $^1$HNMR, GPC, and FTIR Spectra for PEtOx-V3 (Nominal DP-10). Residual ethanol from prior stock solution used to prepare sample for $^1$HNMR.

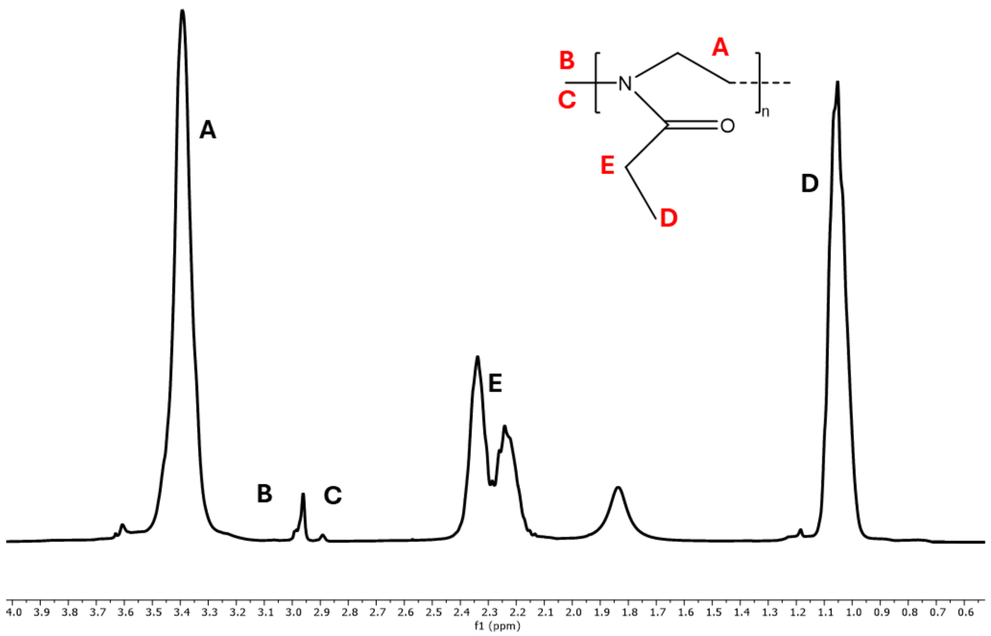


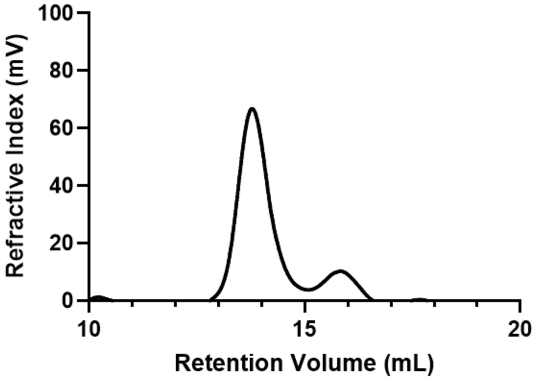


| Peak Label | Peak Range (ppm) | Normalized Area |
|---|---|---|
| A | 3.20-3.57 | 95.63 |
| B+C | 2.86-2.92, 2.94-3.02 | 3 |
| D | 0.96-1.13 | 71.91 |
| E | 2.15-2.44 | 48.10 |

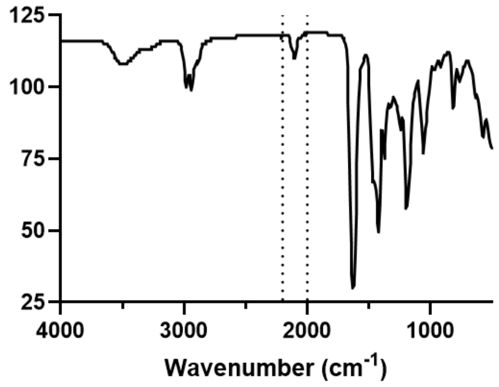


**Supplemental Figure S7:** $^{1}$HNMR, GPC, and FTIR Spectra indicating presence of azide peak from termination for PEtOx-V1 (Nominal DP-20).

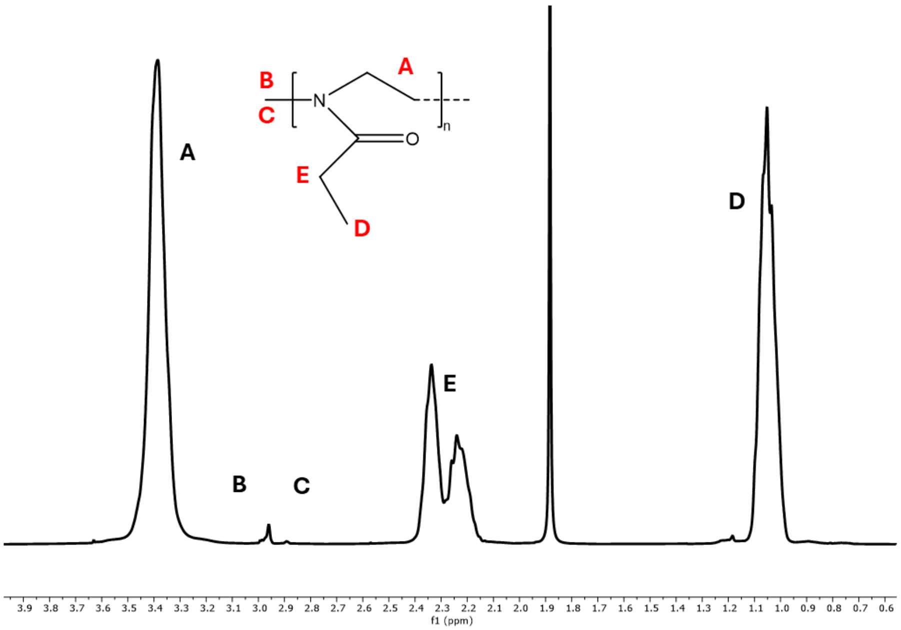


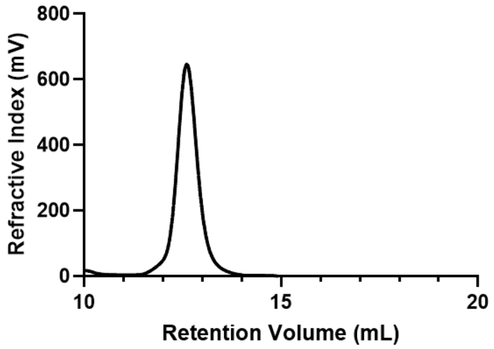


| Peak Label | Peak Range (ppm) | Normalized Area |
| --- | --- | --- |
| A | 3.17-3.57 | 228.37 |
| B+C | 2.86-2.93, 2.93-3.01 | 3 |
| D | 0.95-1.14 | 170.94 |
| E | 2.14-2.42 | 113.96 |

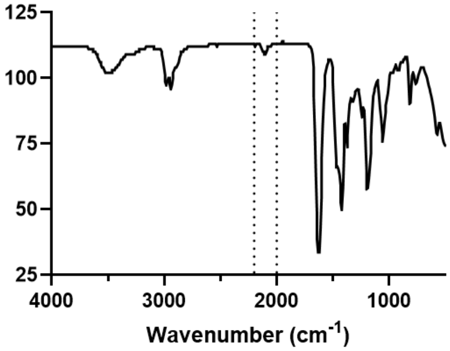


**Supplemental Figure S8:** $^{1}$HNMR, GPC, and FTIR Spectra indicating presence of azide peak from termination for PEtOx-V2 (Nominal DP-40).

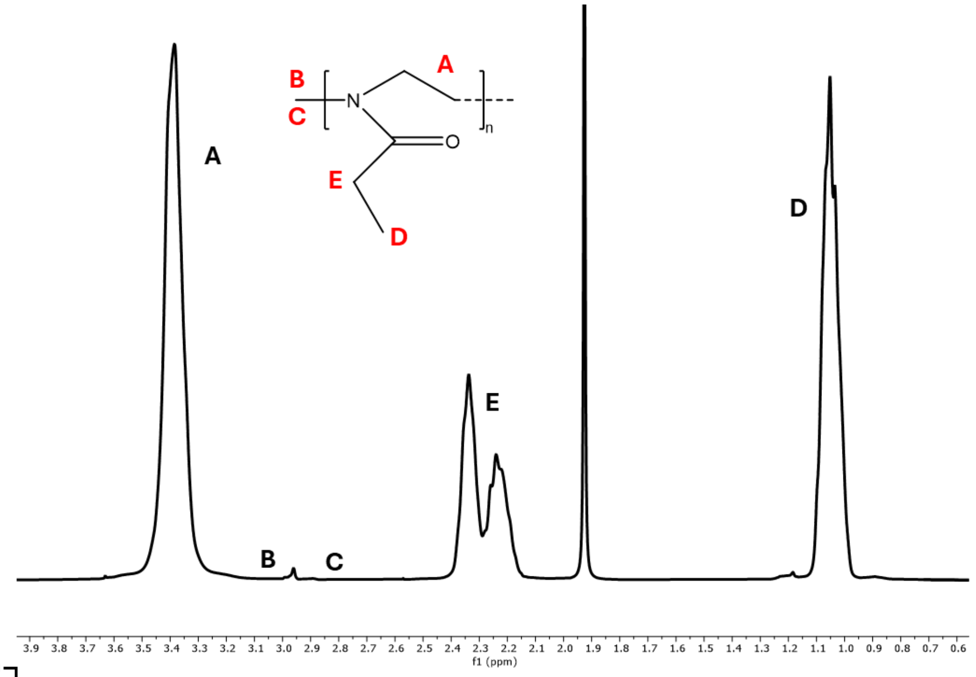


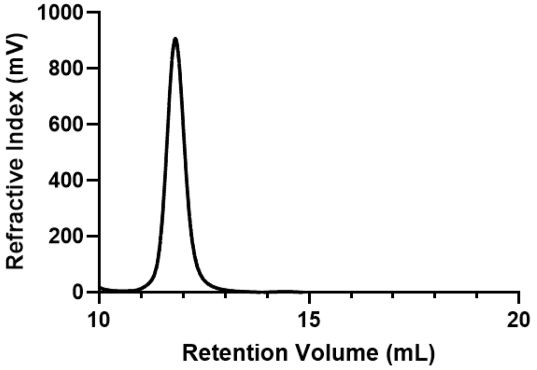


| Peak Label | Peak Range (ppm) | Normalized Area |
|---|---|---|
| A | 3.14-3.61 | 387.96 |
| B+C | 2.87-2.92, 2.93-3.01 | 3 |
| D | 0.96-1.15 | 289.28 |
| E | 2.15-2.43 | 192.99 |

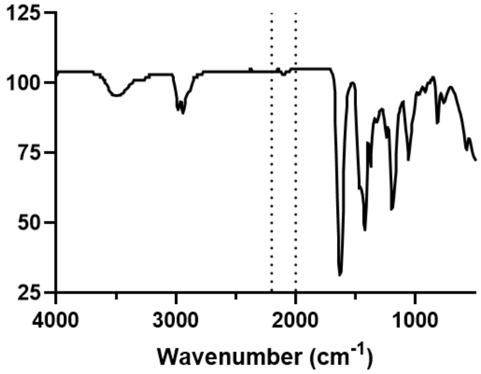


**Supplemental Figure S9:** [1]HNMR, GPC, and FTIR Spectra indicating presence of azide peak from termination for PEtOx-V4 (Nominal DP-80).

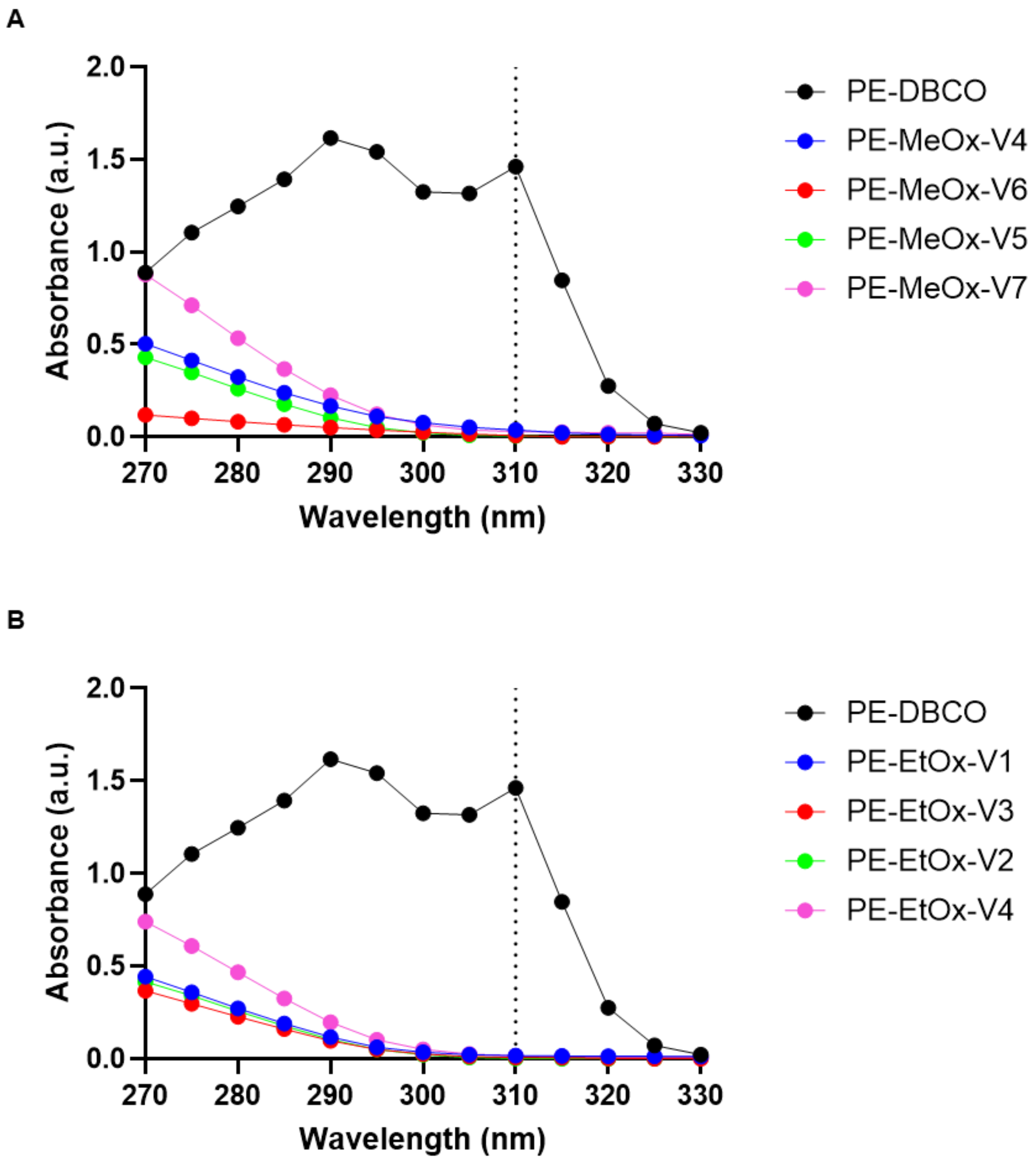


**Supplemental Figure S10:** Decrease of absorbance at 310 nm indicative of SPAAC reaction taking place in **(A)** PMeOx polymers and **(B)** PEtOx polymers.

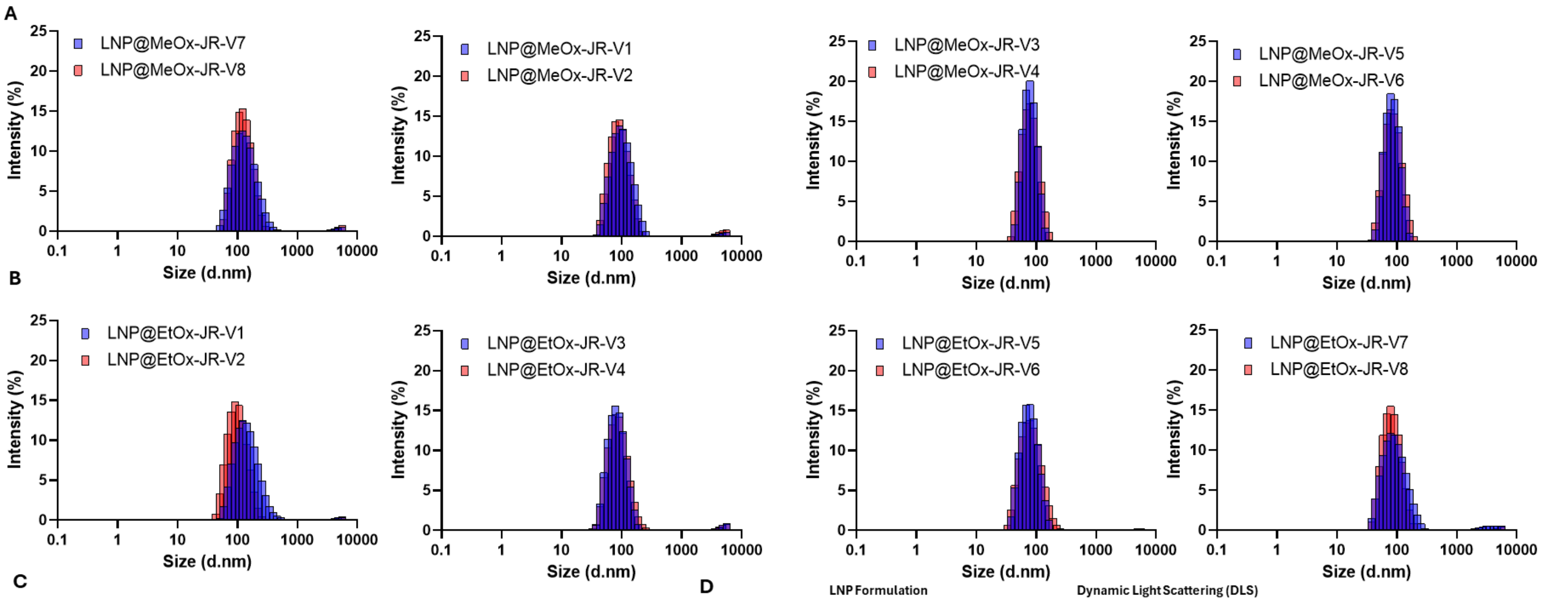


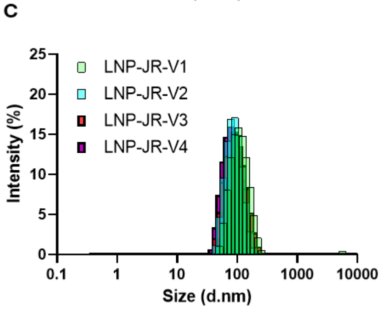


D

| LNP Formulation | | | Dynamic Light Scattering (DLS) | | |
|---|---|---|---|---|---|
| Polymer | Nominal DP | | Intensity Size (nm) | Z-Average Size (nm) | PDI - |
| PEG | MW 2000 g/mol | LNP-JR-V1 | 114.2 | 105.3 | 0.155 |
| | | LNP-JR-V2 | 92.28 | 83.92 | 0.087 |
| | | LNP-JR-V3 | 98.21 | 86.84 | 0.131 |
| | | LNP-JR-V4 | 87.30 | 78.27 | 0.113 |
| PMeOx | DP 10 | LNP@MeOx-V7 | 141.9 | 122.3 | 0.188 |
| | | LNP@MeOx-V8 | 129.7 | 120.2 | 0.172 |
| | DP 20 | LNP@MeOx-V1 | 105.5 | 94.16 | 0.174 |
| | | LNP@MeOx-V2 | 96.19 | 88.68 | 0.182 |
| | DP 40 | LNP@MeOx-V3 | 80.37 | 74.82 | 0.065 |
| | | LNP@MeOx-V4 | 82.10 | 74.44 | 0.101 |
| | DP 80 | LNP@MeOx-V5 | 86.65 | 79.93 | 0.071 |
| | | LNP@MeOx-V6 | 89.08 | 80.27 | 0.099 |
| PEtOx | DP 10 | LNP@EtOx-V1 | 154.6 | 130.0 | 0.182 |
| | | LNP@EtOx-V2 | 105.0 | 95.08 | 0.161 |
| | DP 20 | LNP@EtOx-V3 | 87.46 | 81.88 | 0.182 |
| | | LNP@EtOx-V4 | 92.14 | 83.50 | 0.182 |
| | DP 40 | LNP@EtOx-V5 | 80.46 | 73.33 | 0.130 |
| | | LNP@EtOx-V6 | 86.32 | 75.07 | 0.154 |
| | DP 80 | LNP@EtOx-V7 | 98.48 | 86.81 | 0.210 |
| | | LNP@EtOx-V8 | 86.28 | 78.87 | 0.167 |

**Supplemental Figure S11:** DLS Histograms for **(A)** PMeOx LNPs (increasing DP from 10 (left) to 80 (right) **(A)** PEtOx LNPs (increasing DP from 10 (left) to 80 (right) **(C)** PEG-2000. **(D)** Table of sizes for each LNP formulation.

**Supplemental Table S3:** Individual batch characterization of PEG and POx LNPs prepared for this work

| LNP Formulation | | Dynamic Light Scattering (DLS) | | | Nanoparticle Tracking Analysis (NTA, Zetaview) | | | | | RiboGreen RNA Quantification Assay | |
|---|---|---|---|---|---|---|---|---|---|---|---|
| | | Intensity Size | Z-Average Size | PDI | Mean Size | Median Size | Mode Size | Total Particles | Zeta Potential | Total mRNA Concentration | Encapsulation Efficiency |
| Polymer | Nominal DP | (nm) | (nm) | - | (nm) | (nm) | (nm) | - | (mV) | (ug/mL) | - |
| PEG | MW 2000 g/mol | 114.2 | 105.3 | 0.155 | 91.3 | 81.8 | 78.4 | 7.6E+11 | -5.2 | 237.5 | 93.6% |
| | | 92.28 | 83.92 | 0.087 | 83.8 | 77.5 | 77.4 | 1.5E+12 | -5.20 | 276.4 | 91.6% |
| | | 98.21 | 86.84 | 0.131 | 83.1 | 76.2 | 74.3 | 9.0E+11 | -4.25 | 262.0 | 91.5% |
| | | 87.30 | 78.27 | 0.113 | 81.5 | 74.6 | 73.1 | 9.8E+11 | -7.13 | 232.0 | 91.1% |
| PMeOx | DP 10 | 141.9 | 122.3 | 0.188 | 99.5 | 86.9 | 82.0 | 7.6E+11 | -7.94 | 186.8 | 83.6% |
| | | 129.7 | 120.2 | 0.172 | 99.1 | 88.1 | 83.0 | 7.2E+11 | -10.89 | 220.1 | 86.6% |
| | DP 20 | 105.5 | 94.16 | 0.174 | 85.0 | 76.1 | 73.3 | 5.9E+11 | -4.11 | 218.7 | 89.0% |
| | | 96.19 | 88.68 | 0.182 | 81.2 | 73.5 | 71.7 | 1.3E+12 | -5.68 | 207.1 | 87.1% |
| | DP 40 | 80.37 | 74.82 | 0.065 | 83.0 | 76.7 | 75.5 | 1.1E+12 | -3.12 | 231.4 | 86.1% |
| | | 82.10 | 74.44 | 0.101 | 82.4 | 75.2 | 74.1 | 1.1E+12 | -3.00 | 179.0 | 85.4% |
| | DP 80 | 86.65 | 79.93 | 0.071 | 83.6 | 77.0 | 75.9 | 1.1E+12 | -2.73 | 183.3 | 81.1% |
| | | 89.08 | 80.27 | 0.099 | 84.3 | 78.2 | 77.1 | 1.2E+12 | -4.84 | 185.6 | 88.9% |
| PEtOx | DP 10 | 154.6 | 130.0 | 0.182 | 102.4 | 89.7 | 82.4 | 8.9E+11 | -7.24 | 183.4 | 77.6% |
| | | 105.0 | 95.08 | 0.161 | 85.7 | 77.8 | 75.6 | 8.4E+11 | -12.55 | 192.7 | 77.7% |
| | DP 20 | 87.46 | 81.88 | 0.182 | 79.7 | 71.6 | 70.1 | 1.0E+12 | -3.36 | 246.1 | 84.0% |
| | | 92.14 | 83.50 | 0.182 | 79.8 | 72.5 | 71.6 | 9.6E+11 | -4.14 | 273.2 | 84.0% |
| | DP 40 | 80.46 | 73.33 | 0.130 | 81.0 | 74.4 | 72.7 | 8.8E+11 | -3.82 | 236.2 | 81.5% |
| | | 86.32 | 75.07 | 0.154 | 80.1 | 74.1 | 73.4 | 8.1E+11 | -4.78 | 235.6 | 79.0% |
| | DP 80 | 98.48 | 86.81 | 0.210 | 87.2 | 80.0 | 78.4 | 6.8E+11 | -0.87 | 238.0 | 79.3% |
| | | 86.28 | 78.87 | 0.167 | 84.0 | 78.5 | 78.0 | 2.4E+12 | -5.79 | 168.6 | 74.7% |

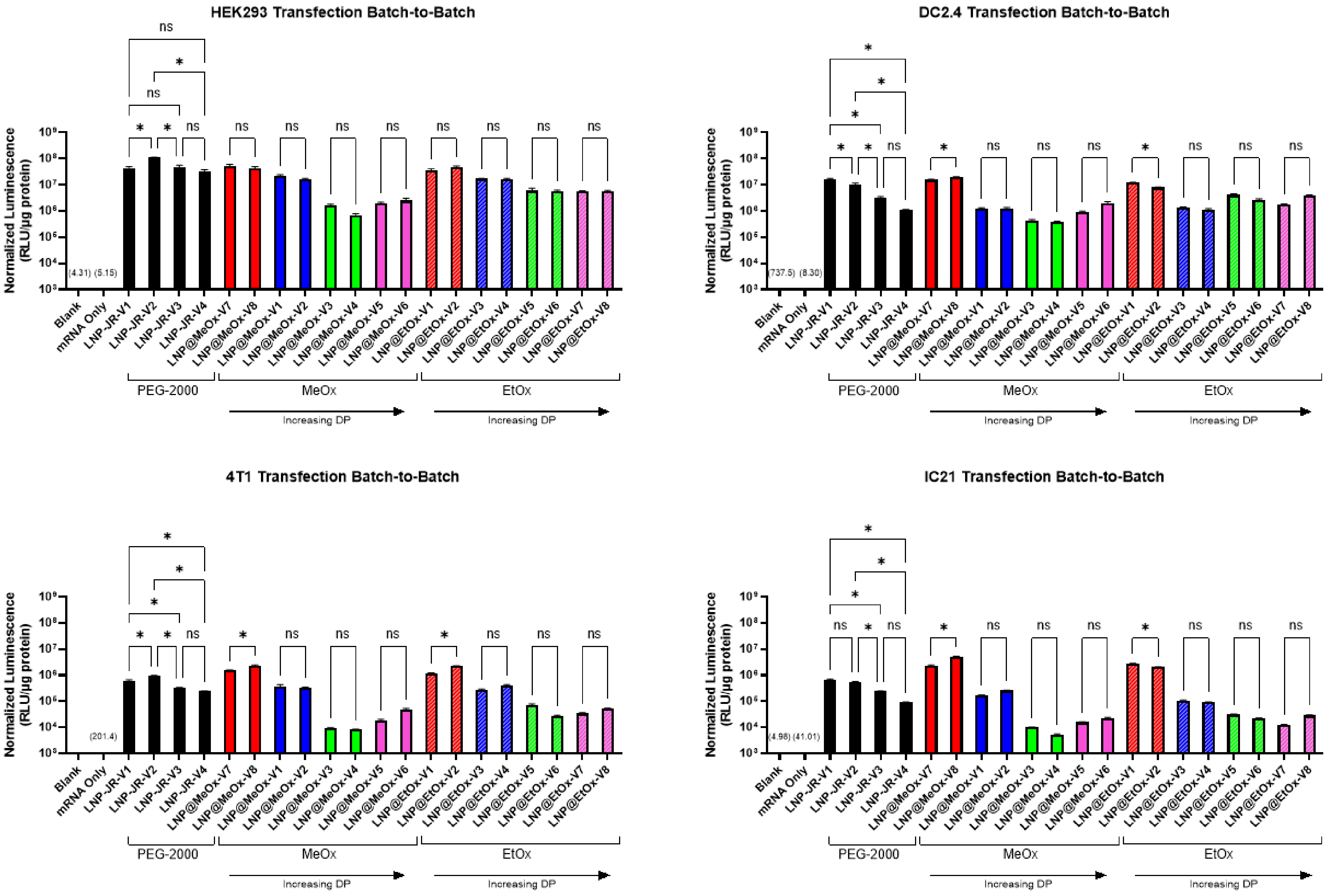


**Supplemental Figure 12:** Batch-to-batch variation in transfection of HEK293, DC 2.4, and 4T1 cell lines. One-Way ANOVA analysis was performed to evaluate batch-to-batch variation of each LNP formulation. Overall α=0.05 with corrected p-values. * indicates significance with adjusted p-value<0.05. Data presented is mean ± SEM with n=6 replicates per batch (Except IC21 LNP@MePx-JR-V1, n=3 due to insufficient material).

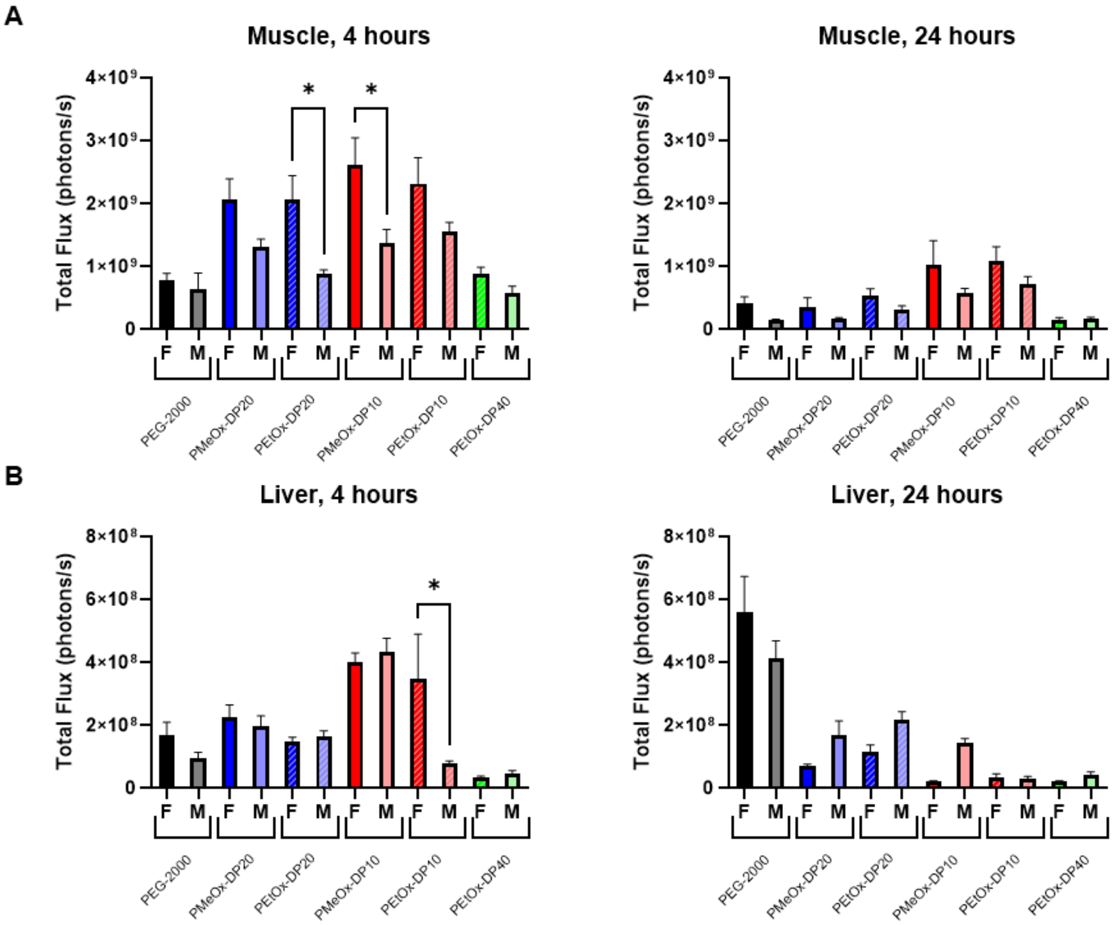


**Supplemental Figure 13:** Comparison of female and male transfection levels from *in vivo* IVIS luminescent imagining. Graphs compare transfection in **(A)** Muscle tissue at 4 and 24 hours and **(B)** Liver at 4 and 24 hours. Intersex variability as evaluated using one way ANOVA analysis with intersex comparisons (α=0.05). N=3 mice/gender/formulation.

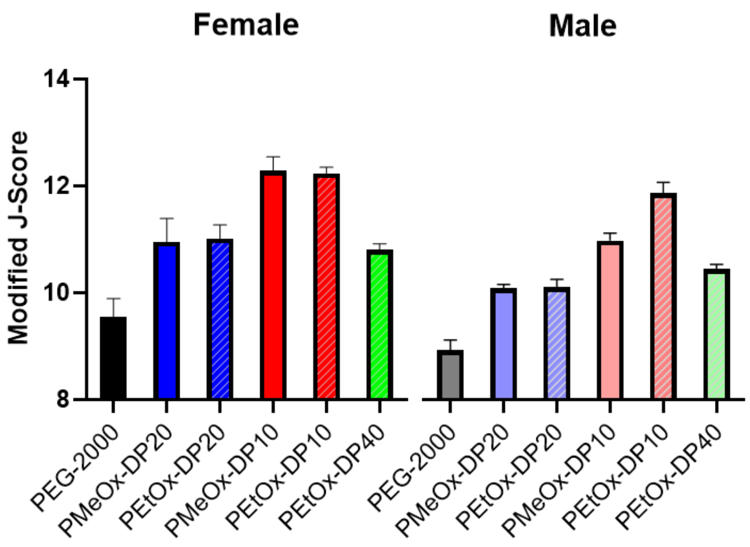


**Supplemental Figure 14:** Modified J-Scores which account for expression intensity level differences between formulations calculated using **Equation 3**.

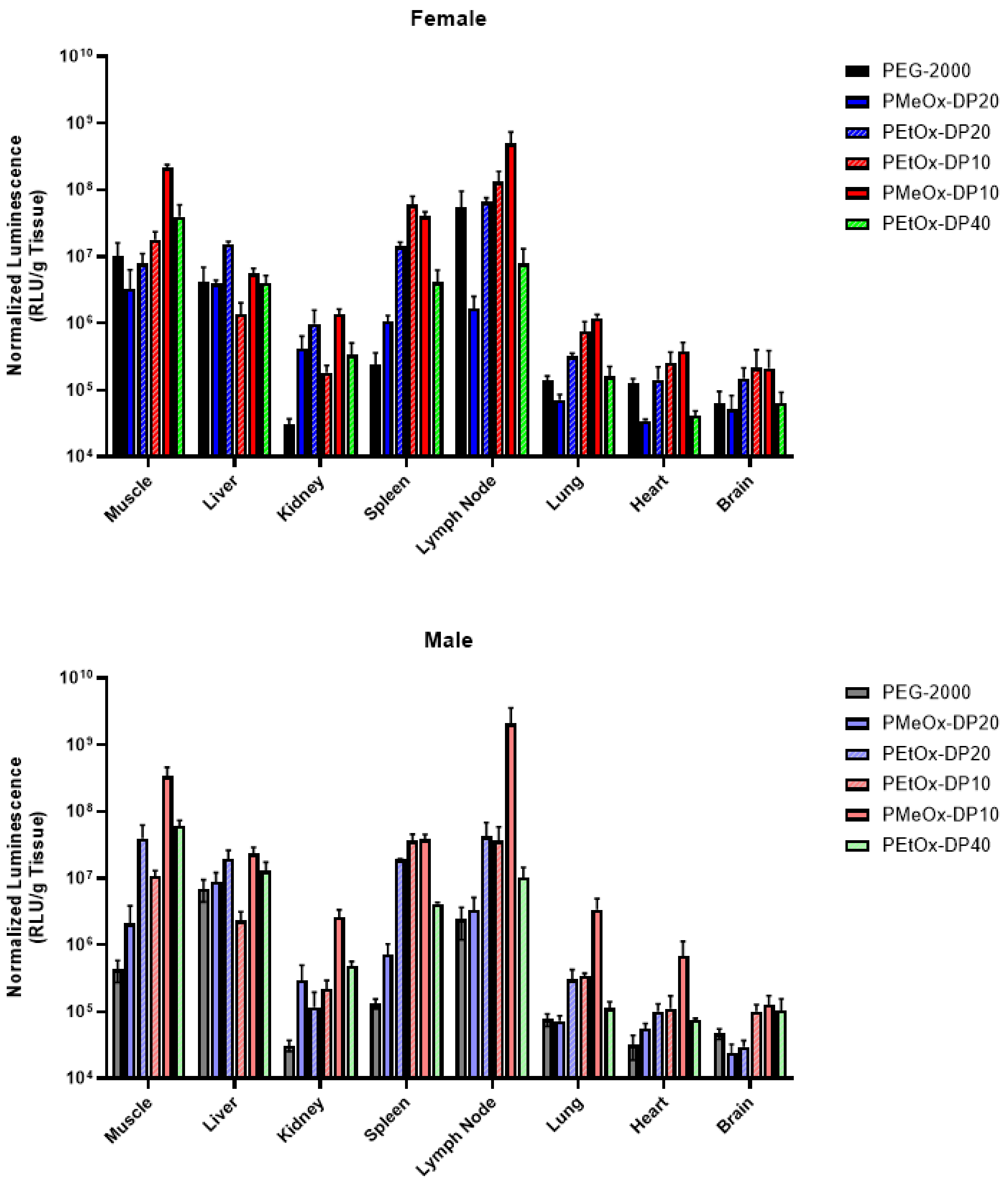


**Supplemental Figure S15:** Luminescence per gram of tissue in various isolated tissues 24 hours after LNP injection in both male and female mice.

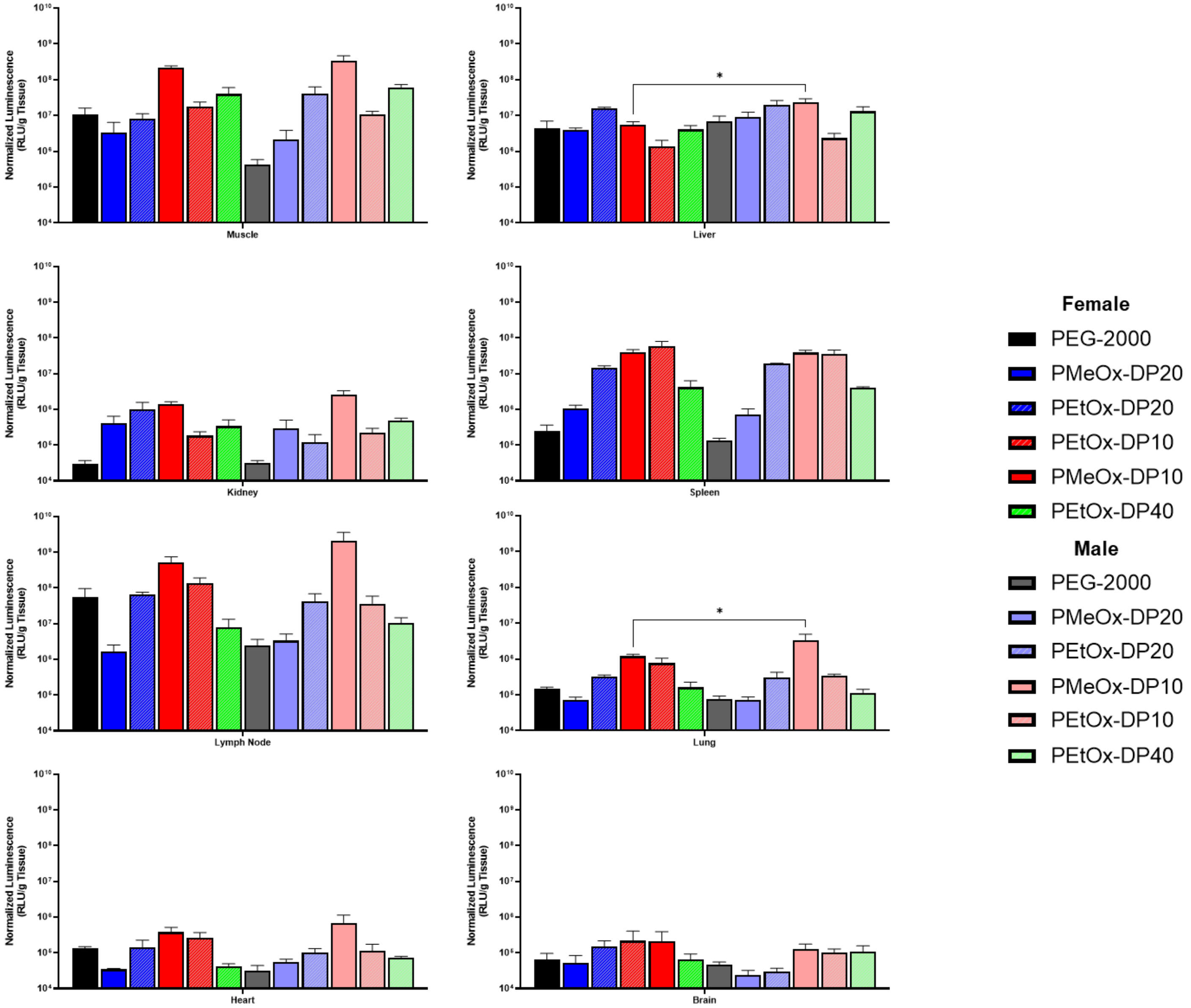


**Supplemental Figure S16:** Comparing intersex differences of each tissue as a function of LNP formulation.
